# Single-molecule landscape of DNA replication pausing

**DOI:** 10.1101/2025.08.14.670160

**Authors:** Sathish Thiyagarajan, Anna M. Rogers, Carolin A. Müller, Conrad A. Nieduszynski

## Abstract

A pause in DNA synthesis that occurs when the replisome encounters an obstacle could lead to genome instability. Although important, systematic identification of replication pause sites is challenging due to their low frequency and delocalized nature. Here we present the first single-molecule identification of sites of replisome perturbation across a eukaryotic genome using long-read nanopore sequencing. For each single-molecule replication pause we determine the direction of replication, leading/lagging-strand identity, location and approximate duration, and whether the replisome resumed synthesis. Although pauses are largely diffuse over the genome, they are significantly enriched over transcribed features and correlate with transcription and R-loop levels. Transcription-replication conflicts are more numerous when head-on than co-directional. Finally, we identified genomic loci with a strong bias towards leading over lagging strand pauses, consistent with uncoupling of the helicase from polymerase epsilon. Our data support helicase-polymerase uncoupling resulting from replication pausing as the molecular trigger behind epigenetic switching.

**Graphical abstract:** 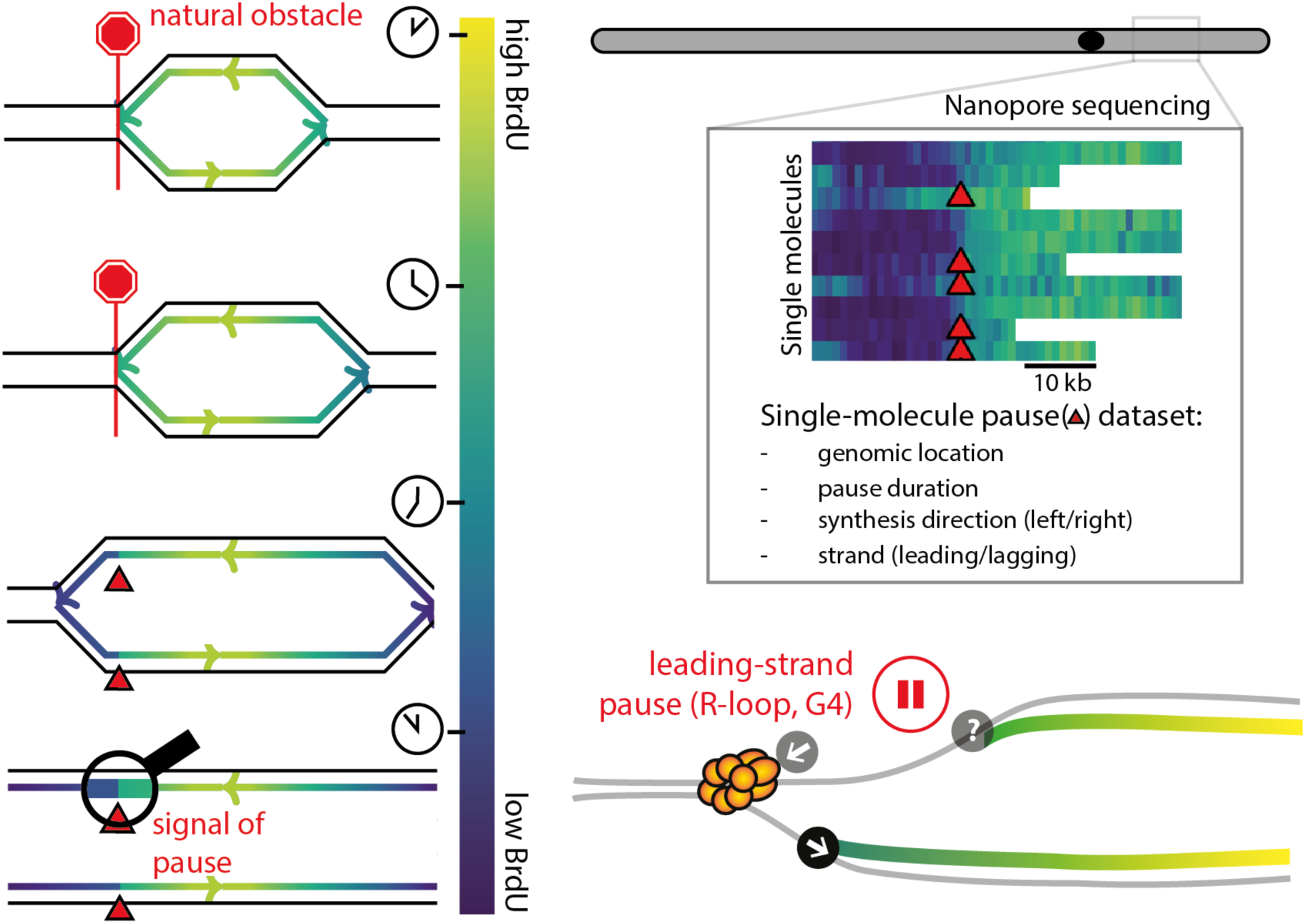

## Introduction

Faithful DNA replication is essential for life, with errors giving rise to genetic variation that can be beneficial or deleterious. Although such errors are rare, their impact can be dramatic, including diseases such as cancer^1^. Therefore, cells tightly regulate the dynamics of DNA replication at the level of initiation, termination, and movement of the replisome, the DNA-synthesizing protein complex, at the replication fork. Challenges to fork progression pose a particular risk to genome stability, as a pause increases the likelihood of single-stranded DNA and breakage^2^. However, currently available techniques have not been able to simultaneously resolve the location, frequency, and duration of replisome pausing.

DNA replication must overcome a number of challenges encountered on the DNA template, such as non-canonical secondary structures^3,4^, lesions^5^, transcription^6^, and bound proteins^7^. These challenges are stochastic as they arise intermittently^8–10^ and are distributed widely across the genome^11–15^. Thus, cells have evolved sophisticated mechanisms to minimize and overcome problems. For instance, cells coordinate transcription and replication^16^ and accessory helicases can be recruited to the replisome to overcome obstacles^17^. Overall, the replisome achieves a uniform average synthesis rate^18–20^, but problems still arise.

In prior studies, detection of pauses in replication progression was limited due to the reliance on ensemble population data^3–7,12–14,19,21^ or ensembles of single molecules^20^. A consequence of averaging across millions of cells is the enrichment of loci where pauses are tightly focused spatially on the DNA and the loss of loci with dispersed pauses in the noise. In addition, at any locus, long-lasting pauses in a minority of cells would either be lost or would produce a similar signal as shorter pauses that affect many cells. Furthermore, at most loci, cell-to-cell variation in both replisome direction and the leading or lagging strand identity at a paused site may also be lost. In summary, currently we cannot quantify the fraction of replisomes that pause at a particular location or the associated pause duration, therefore the landscape of problems in DNA replication is unclear.

Here we present the first whole genome single molecule identification of sites of replisome perturbation. We utilized temporal variation in the level of cellular incorporation of a synthetic nucleoside analogue (Bromodeoxyuridine or BrdU), which we detected via direct nanopore sequencing. This allowed measurement of *in vivo* DNA replication kinetics and determination of sites of replisome pausing. Our single-molecule approach has equal sensitivity to pause sites irrespective of whether they are frequent or rare in a population of cells, or whether they are tightly focused spatially on the DNA across the population or are spatially dispersed. We measured the location and duration of replisome pauses throughout the genome and related them to the landscape of transcription and R-loops—non-canonical DNA-RNA hybrids, a subset of which are genotoxic^22^. Although enriched near transcribed regions, most pauses were diffusely distributed across the non-repetitive genome. Finally, we identified strand-specific replication pauses that are consistent with helicase-polymerase uncoupling and which may lead to epigenetic switching.

## Results

### Distinct leading and lagging strand pauses sites on single molecules at the rRFB

As a first step to replisome pause detection, we inferred replication time from individual molecules using a BrdU-aware nanopore sequencing workflow. Work from us and others has shown that when budding yeast (expressing a nucleoside transporter and thymidine kinase) are exposed to a constant BrdU concentration, the replisome incorporates progressively lower BrdU levels in synthesized DNA versus time in S phase^23,24^. This is presumably due to endogenous thymidine synthesis^25^. In this way, local BrdU density on single molecules becomes a proxy for replication time.

This technique allows us to capture replication events on a single-molecule basis (equivalent to single-cell), with high spatial and strand resolution. Peaks, smooth gradients, and abrupt steps in BrdU density are interpreted as initiations, normal replisome tracks and sites of replication pausing respectively (Fig. 1A, Videos 1 and 2)^23^. We interpret a sudden BrdU decline as a paused site as it implies an abrupt increase in the local time of DNA synthesis along a nascent molecule (Video 2). Using this technique, we previously demonstrated detection of initiation sites and performed a proof-of-principle low spatial-resolution analysis of pausing at the well-characterized unidirectional replication fork barrier (RFB) in the ribosomal DNA (rDNA) repeats. Here, we have generated an ∼8x larger whole genome dataset of 6.9 million molecules (Figs. 1A, Fig. S1 and *Methods*), used improved analogue detection algorithms^26^ and a novel computational framework to identify rare, but physiologically important, pauses in DNA synthesis.

**Figure 1.**
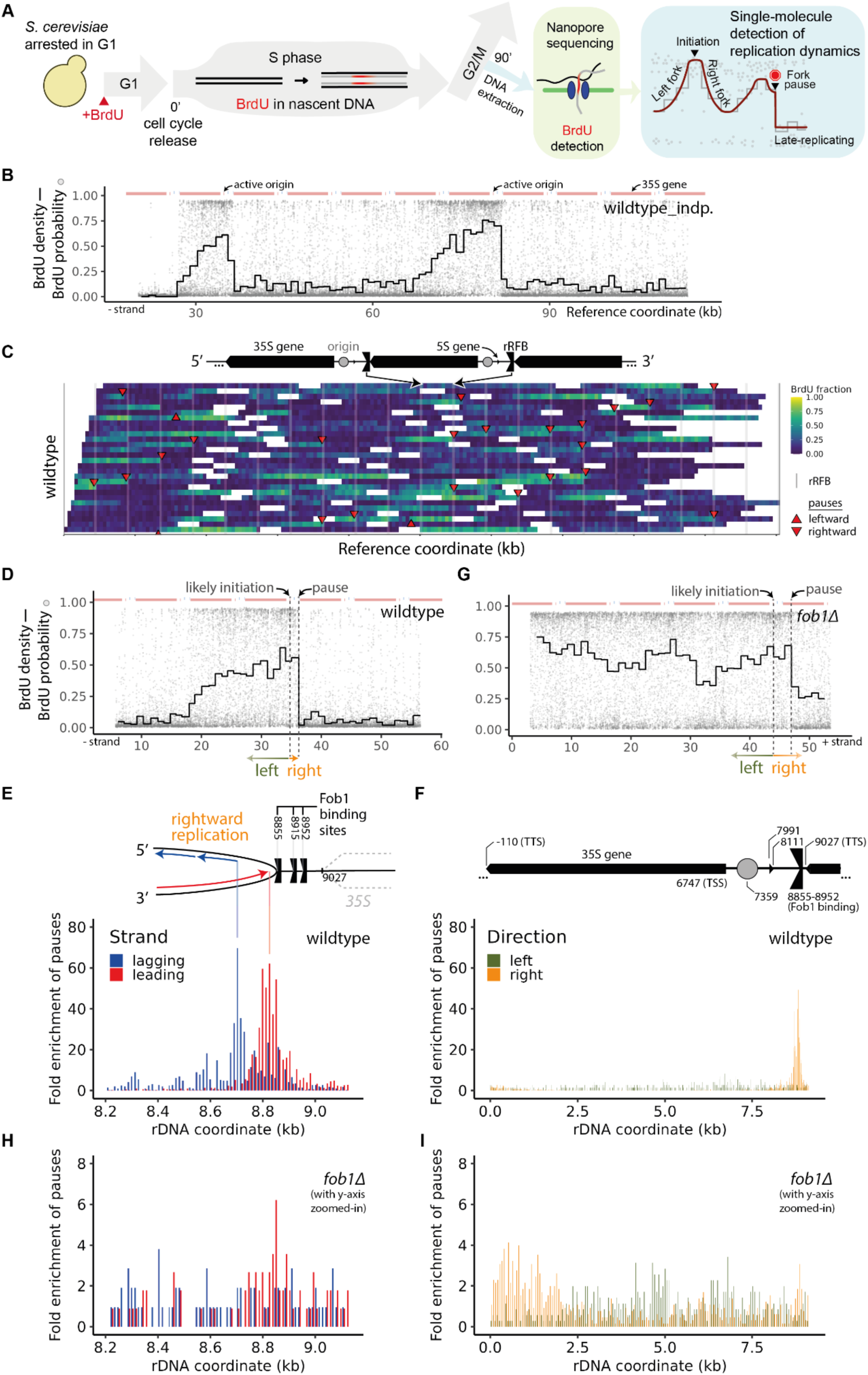
Single molecule analysis reveals distinct spatial patterns of leading and lagging strand pauses in the ribosomal DNA. **(A)** Schematic of the protocol to record replication dynamics using BrdU density in *S. cerevisiae*. Synchronized cells incorporate BrdU in DNA during S-phase at decreasing levels over time^23^. We nanopore sequenced DNA and measured BrdU substitution probabilities per thymidine per molecule. On single-molecules, we detected peaks, gradients, and sharp declines in BrdU density and labelled them as initiations, replisome tracks, and replication pauses respectively. **(B)** Single-molecule BrdU density versus genomic coordinate of a ∼100 kb molecule with two initiations in the rDNA. Grey points: BrdU substitution probability per individual thymidine. Black: average BrdU substitution rate in non-overlapping windows of 300 thymidines each. Top: genomic annotations along the rDNA. **(C)** Top: Schematic of rDNA repeats. Bottom: Heatmaps of BrdU density of a randomly selected subset of nascent molecules aligned to the rDNA in wildtype (*n* = 100 molecules). Sharp declines in BrdU density are detected as replisome pauses (arrowheads; down/up refer to rightward/leftward pauses respectively); these are almost all rightward pauses and cluster at the rRFBs (vertical grey lines). Heatmaps show BrdU density per molecule per 1 kb-sized windows along the reference. Molecules stacked horizontally and vertically such that neighbouring molecules on same row are separated by at least 5 kb (white). **(D)** Single-molecule BrdU density versus genomic coordinate of a molecule with a detected replisome pause. Likely initiation site and movement of leftward and rightward replisomes are highlighted. **(E)** Top: Schematic of a rightward replisome approaching the rRFB. Bottom: Fold enrichment of pauses contributed by leading and lagging-strand syntheses versus mapped rDNA coordinate near the rRFB (*n* = 3041 pauses). Data are collapsed onto one rDNA repeat. Wildtype leading and lagging strand syntheses arrest 33 bp and 150 bp behind the barrier respectively (modal values, vertical lines). **(F)** Top: Schematic of rDNA repeat. Bottom: as for **(E)** but zoomed out over the whole rDNA repeat and coloured by replisome direction rather than leading or lagging strand identity. **(G)** Single-molecule BrdU density versus genomic coordinate for a molecule with a detected replisome pause from a *fob1Δ* cell. **(H)** As for **(E)** but in a *fob1Δ* cell (*n* = 148 pauses). **(I)** Same as **(F)** but for *fob1Δ* cells.

We first analysed BrdU density around the rDNA unidirectional fork barrier. The rDNA region consists of 150-200 tandem repeats (each ∼9 kb) at one genomic locus, each of which includes a DNA replication initiation site, RNA polymerase transcribed units, and an RFB. The RFB limits transcription-replication conflict by pausing rightward replisomes (opposite to the direction of 35S transcription) and depends upon the protein Fob1 binding at three close, distinct sites^27–29^. However, not all 35S genes and initiation sites are active and estimates of the fraction vary^9,29–32^. The single molecule BrdU density peaks within the rDNA are highly asymmetric, with a significantly sharper rightward gradient than leftward (Fig. 1B). The pattern is indicative of replication initiation, followed by a rightward fork pause, while the sister leftward fork progressed uniformly. To measure pauses quantitatively, we detected locations of sudden BrdU density changes along each molecule (see *Methods*). We observed replication initiation only in a subset of repeats, consistent with most of the rDNA being passively replicated^29,32^, although we cannot identify origin firing in later S phase when BrdU incorporation is already low (Fig. 1C).

Unlike short-read sequencing, reads of ten kilobases and above allow us to see spatial patterns of replication across multiple rDNA repeats from a single cell. Multiple initiations and pauses are seen in the same molecule (Fig. 1B). The rDNA repeats to the right of the pauses in Figs. 1B, D are presumably passively replicated by leftward replisomes which arrive at the ‘active’ barrier to complete replication and terminate with the paused fork. Assuming a mean fork speed of 2 kb min^-118,19,33^, a replisome should take ∼4.5 mins to synthesize one rDNA repeat. The leftward replisomes of Figs. 1B, D that start moving at the time of high BrdU replicate ∼1-2 rDNA repeats before the levels drop significantly. This suggests that at least ∼5-10 minutes elapse before the rescue during low BrdU of the arrested sister rightward forks (Figs. 2A, B).

**Figure 2.**
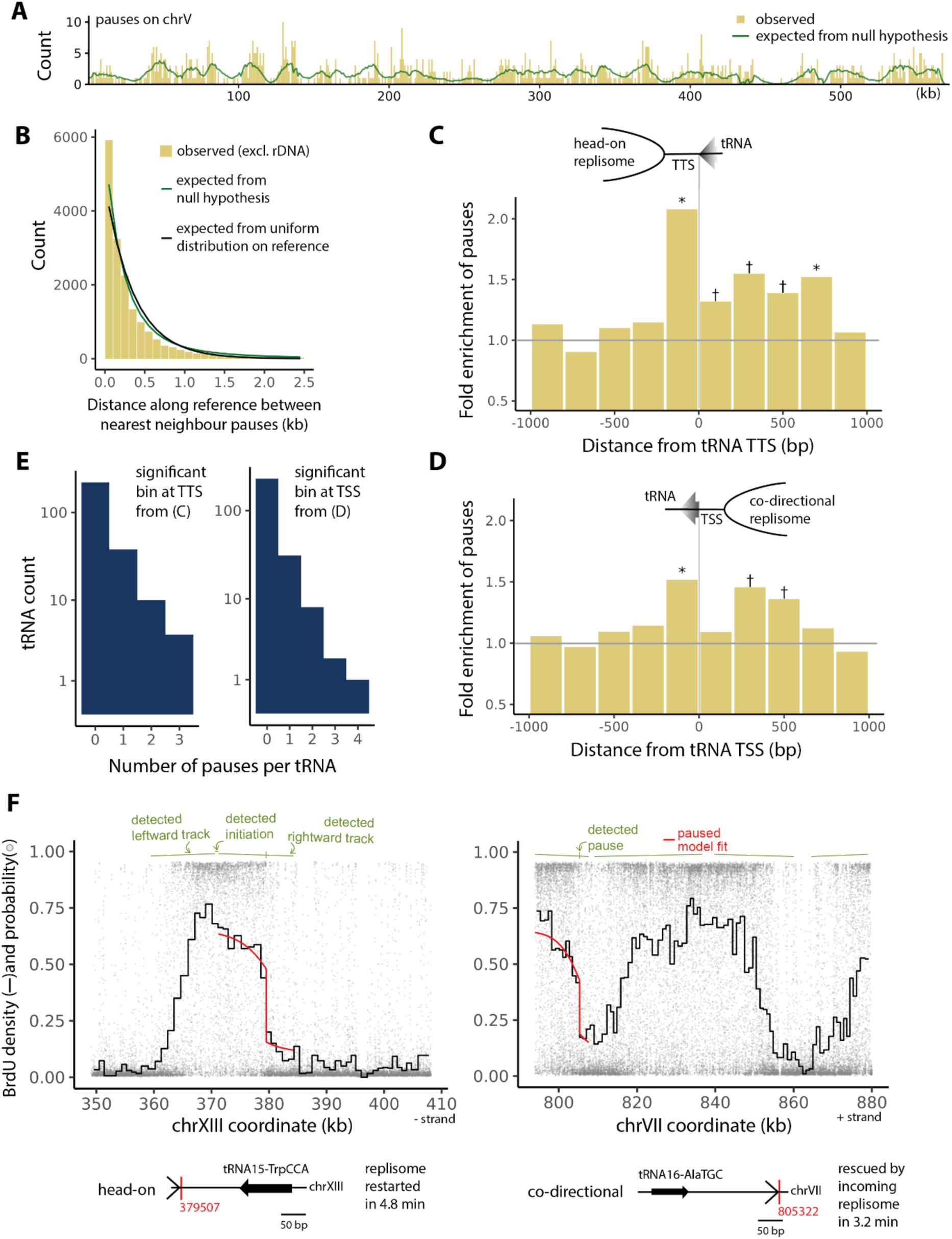
Replisome pauses are enriched in proximity to tRNAs. **(A)** Expected and observed count of single-molecule replisome pauses versus their mapped location along chromosome V (*n* = 953 pauses; for other chromosomes see Supplementary Fig. S4). **(B)** Histogram of expected and observed nearest-neighbour distances between replisome pauses along reference genome, excluding rDNA (*n* = 16846 distances). Black: expectation from spatially uniform distribution of pauses. **(C, D)** Ratio of observed to expected head-on **(C)** and co-directional **(D)** pause counts versus distance of pause relative to nearest tRNA transcription termination **(C)** and start **(D)** sites (*n* = 446 **(C)** and 458 **(D)** pauses, * and † indicate > 3 s.d. or > 1.65 s.d. respectively of enrichment). **(E)** Histogram of pause counts per tRNA gene in the central bins of **(C)** and **(D)** (*n* = 70 and 57 pauses respectively). **(F)** Top: single-molecule BrdU density versus genomic coordinate of two molecules showing pausing on head-on (left) and co-directional (right) replisome tracks relative to tRNA transcription. Red: best-fit model with pause. Green: replisome track annotations. Bottom: diagram of pause location and replisome orientation relative to tRNA and nature of replication rescue of corresponding events. Genomic coordinates from W303 reference^59^.

We plotted the location of detected pause sites on a single rDNA repeat (Figs. 1E, F, S2A). This approach allowed us to visualise the spatial distribution of pauses relative to the major rDNA features. As anticipated, we observed that fork pause sites were highly clustered at the fork barrier, predominantly on rightward replisomes and at a level ∼50x that expected from a uniform distribution across the repeat (Figs. 1E, F, Table S2). Within the distribution there are two distinct peaks of pausing proximal to the barrier (see below; Fig. 1F). Moving from left to right on molecules across pause sites, we see a gradual increase, a sharp drop at the pause, and then a gradual increase in BrdU density (Fig. S2B). The two gradual increases are signatures of leftward replisomes—the sister of the paused replisome moving away from their common initiation site, and the rescuing replisome arriving at the pause site respectively. Additionally, we noted a minority of replisome tracks that lack a sharp drop off over the RFB, consistent with forks not being arrested (Fig. S2C). Multiple pauses along the same molecule were in register with the repeat length (Fig. S2D); we have fewer molecules with widely separated pauses because longer molecules are rarer in the dataset (Fig. S1B).

We separated pauses into those observed from leading and lagging strand synthesis. This was determined from replisome direction and whether the nascent strand was the top or the bottom strand (top of Fig. 1E). Leading and lagging strand pauses were 33 bp and 150 bp behind the first Fob1 binding site, respectively (modal values, *n* = 2145 and 2048 pauses respectively) (Figs. 1E, S2E). We measured similar pause locations in our independent dataset (Fig. S2F). The leading strand pause is more focused and barrier-proximal, with the separation from the barrier similar to the ∼20-40 bp footprint of a CMG helicase^34^, and the separation between the strand pauses is about the modal length of an Okazaki fragment^35^. Thus, we quantify pauses at the ribosomal replication fork barrier with high strand-specific spatial resolution.

### Transcription-replication conflict observed in the absence of the RFB

To confirm that the rDNA fork barrier is primarily responsible for the fork pauses we detected at the rDNA locus, we measured BrdU patterns in *fob1Δ* cells that lack the barrier-creating Fob1 protein^27,28^ (Table S1). We used the same pause-detection approach as wildtype. In an example single molecule (Fig. 1G), we detect a pause, but it is not proximal to the rRFB nor is it associated with a characteristic asymmetric BrdU peak seen at the wildtype rRFB (Figs. 1B, D). We plotted all detected pauses onto one rDNA repeat for visualisation (Figs. 1H, I). Pauses were quite diffuse—the strong wildtype pause signal at the rRFB was absent. BrdU step sizes were smaller at the pause, suggesting a smaller pause duration (Fig. S2B). Without an operational fork barrier, we expect the rightward replisome to synthesize more DNA during the time of high BrdU compared to wild type. Therefore, we expected and observed more BrdU-positive regions in *fob1Δ* (Figs. 1C, 1G and S2G, S2H).

In the *fob1Δ* dataset, we measured an enrichment of rightward pauses starting at the transcription termination site of the highly expressed 35S rRNA gene and extending ∼ 2 kb into the gene body (maximum enrichment of ∼4.1x, Fig. 1I). As the termination site is just ∼170 bp beyond the barrier, unhindered rightward replisomes would first experience conflicts with 35S transcription here in a head-on manner (see schematics of Figs. 1E, F). We also measured an enrichment of leftward or co-directional pauses extending from the transcription start site and a similar length of ∼2 kb into the gene body (peak enrichment of ∼3.4x, Fig. 1I), but these are fewer in number—there were 85% more rightward than leftward pauses (Table S2). In summary, in the absence of a functional RFB, we observe what are likely transcription-replication conflicts, with head-on conflicts being more numerous than co-directional.

### Detection of pauses in the non-repetitive genome

Next, we analysed the non-repetitive portion of the *S. cerevisiae* genome. While we expect precipitous drops in BrdU density at pause sites like those at the rDNA, we did not assume the tight spatial clustering seen at the rDNA, nor that replisomes would pauses for such extended timespans (briefer replisome pauses would give smaller drops in BrdU density). Therefore, we used a more sophisticated approach to identify sharp discontinuities along individual replisome tracks. We first established that the BrdU decline followed a characteristic smooth shape across the replication track, and using this pattern as a reference identified the sites along a minority of tracks that had sharp discontinuities in BrdU density (Fig. S3; also see *Methods*). This approach also allows us to infer an approximate pause duration from the step size, assuming a synthesis rate of 2 kb min^-118^.

We determined: (1) the location, duration, and replisome direction of significant individual pauses, limited to 0 or 1 pause per replisome track, and (2) a null hypothesis of expected pause count (with respect to fork direction) along the genome (Figs. 2A, S4, S5A). Overall, we examined 330,000 replisome tracks and identified 17,000 pauses: a pause rate of 1 in ∼20 tracks, which implies 5% of replisomes pause per cell per S-phase (Figs. 2A, S4, Table S3). Pauses had a duration of 4.4 ± 1.6 min (mean ± s.d., Fig. S5A). Nevertheless, the smaller drop in BrdU density at shorter pauses makes them harder to distinguish from experimental noise and more likely to be removed from our dataset by the filters we applied to limit false positives (see *Methods* and Supplementary note 2). In an independent dataset, the pause counts per sequenced molecule and the pause durations were similar (Table S3; Fig. S5A), indicating that we are sampling a similar pausing landscape in both datasets. Thus, we can detect pauses genome-wide and not just at strong pausing sites like the rRFB.

### Most replisome pausing is diffusely distributed across the non-repetitive genome

Some regions of the genome are reported to be hard to replicate and by implication should consistently affect replisomes across many cells. This implies that other genomic regions should be easier to replicate, but a quantitative measurement of such a landscape of ‘replication difficulty’ or ‘likelihood of replisome pausing’ along the genome is lacking. To characterize the pause landscape, we placed our single-molecule replisome pause sites on the reference genome and calculated the observed and expected distance between nearest neighbours (see Supplementary note 1). As this is an analysis performed across a population of cells, neighbouring events are likely independent i.e. arise from pauses on replisomes in different cells. This analysis indicates only a minor enrichment (observed to expected ratio of ∼1.2) of replisome pauses with a close neighbour (within 100 bp); most replisome pauses are diffusely distributed across the genome (Fig. 2B). We obtained a similar result with our independent dataset after accounting for the smaller dataset size (Fig. S5B and Supplementary note 1). We conclude that the pause landscape is flatter than previously assumed, with few discrete difficult to replicate loci that pause replication in a high proportion of cells. Most pause events are infrequent and distributed across the genome.

### Quantifying replisome pauses at tRNA genes

Next, we explored replisome pauses at the RNA Pol III-transcribed tRNA genes, where replication fork pauses had been reported previously using population methods^36,37^. Our single-molecule approach and sampling across the genome offers unique insights compared to previous population level analyses: we classify each individual pause as head-on or co-directional with tRNA transcription (as opposed to reliance on ensemble fork direction), we quantify how likely it is for a replisome to pause at tRNA genes versus elsewhere on the genome using our expected pause counts, and visualize the replication history for each event. Thus, we measured the fold enrichment of pauses versus distance from the nearest tRNA transcription start/termination sites (TSS/TTS) in aggregate.

We observed pause enrichment near the tRNA genes (Figs. 2C-F, S5C, D). Head-on and co-directional conflicts were enriched ∼2 fold and ∼1.5 fold respectively at the genes compared to a random genomic locus (central bins of Figs. 2C, D). The distribution of number of pauses per tRNA gene is a rapidly decreasing curve (not bimodal) (Fig. 2E) indicating that tRNA genes do not fall into two distinct groups: those that cause pauses and those that do not. Therefore, pauses are not driven by a minority of tRNAs that are highly problematic compared with a random genomic locus, but by a increased propensity to pausing at a majority of tRNAs.

Two example single molecules, with pauses at tRNAs, demonstrate the diversity of replication events that are detected (Fig. 2F). An example head-on transcription-replication conflict is resolved in ∼5 min and synthesis at the fork restarts. Whereas in a second example, a co-direction transcription-replication conflict was rescued by an incoming replisome in ∼3 min. The other replisome tracks on these molecules appeared to be normal. Thus, we quantified the levels of replication pausing at Pol III-transcribed regions (compared with a random location on the genome) and determine the replication history associated with each pause.

### Replisome pauses are enriched at either end of RNAPII-transcribed regions

Previous studies show replisomes pause at some highly-transcribed RNAPII-genes^6^, so we sought to quantify the pause landscape here. Transcribed genes were divided into quartiles based on their level of transcription (population-level RNAPII occupancy during S phase)^38^ and replisome pauses counted relative to expectation. We separated pauses with respect to direction of encounter (head-on and co-directional) and proximity to the transcription start and termination sites (TSS/TTS, see *Methods*). At transcription start sites, we saw that pauses were significantly enriched, with both relative directions being approximately an equal challenge (Figs. 3A, S6A, B). When split into quartiles, pause enrichment correlated with RNAPII occupancy on replisomes of either relative direction. On average, replisomes were ∼20% more likely to pause over the start sites of the top quartile compared to a random genomic locus.

**Figure 3.**
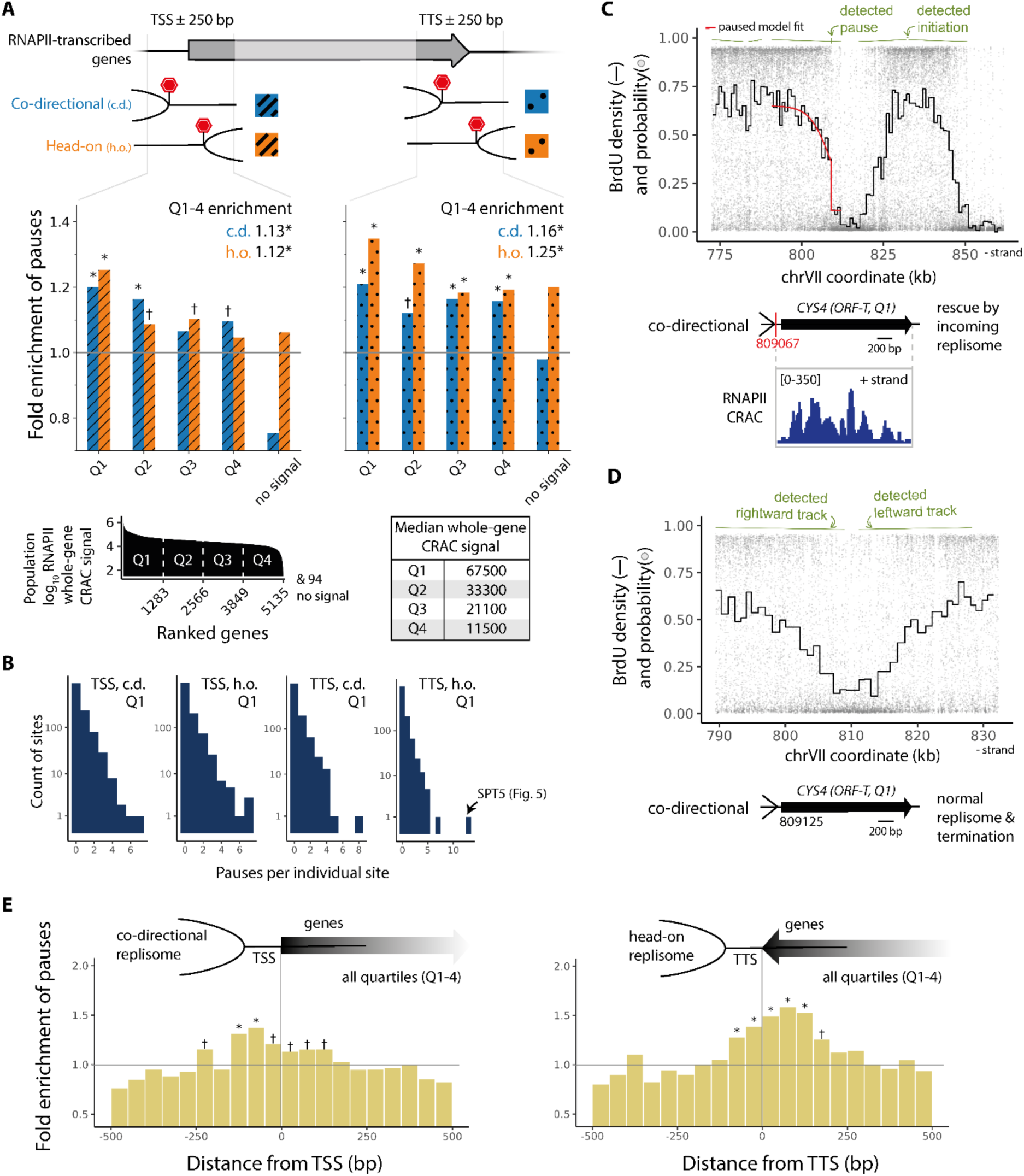
Replisome pauses are enriched near the start and termination sites of RNAPII-transcribed genes and enrichment correlates with RNAPII amount. **(A)** Top: Ratio of observed to expected pause count versus quartiles of total amount of RNAPII occupancy^38^ per transcribed gene at transcription start and termination sites (TSS and TTS respectively). Pauses counted in 500 bp regions centred at TSS/TTS and classified as head-on or co-directional with transcription (see *Methods*). Inset: fold enrichment over all quartiles (see Fig. S6A for *n* values of pauses). Bottom left: Whole-gene RNAPII signal versus genes ranked in descending order. Bottom right: Table of median whole-gene RNAPII signal in each quartile. **(B)** Histogram of pause counts per gene in the four highly transcribed (Q1) bins of **(A)** corresponding to either orientation and TSS/TTS (*n* = 539, 516, 459, and 504 pauses from left to right). **(C, D)** Two example single-molecule BrdU density profiles of molecules displaying a co-directional replisome track over the *CYS4* gene, one with a pause ∼50 bp upstream of its transcription start site that is rescued by an incoming replisome **(C)** and one without any pauses **(D)**. Red: best-fit model with pause. Green: replisome track annotations. Bottom of **(C)**: line diagram of pause location and replisome orientation relative to transcribed gene (ORF-T) and nature of replication rescue, and RNAPII occupancy signal over the gene. **(E)** Ratio of observed to expected count versus relative distance from nearest TSS (left) or TTS (right) of co-directional (left) or head-on (right) pauses in 50 bp bins (all quartiles of transcription; *n* = 2958 and 3038 pauses respectively). Distance measured parallel or anti-parallel to transcription. Throughout, * and † indicate > 3 s.d. or > 1.65 s.d. respectively of enrichment. Genomic coordinates from W303 reference^59^.

At transcription termination sites, differences emerged, although pauses of both relative directions were significantly enriched overall. Head-on pauses were more frequent, and their enrichment increased with greater RNAPII occupancy, whereas co-directional enrichment was relatively RNAPII occupancy-independent (Figs. 3A, S6A, B). Head-on replisomes were ∼35% more likely to pause at the termination site of a top quartile gene than at a random genomic locus; this was ∼20% for co-directional replisomes. Therefore, although replisome pauses are significantly enriched over transcribed regions, most replisomes pass through such regions without problems – consistent with known pathways that mediate and minimise transcription-replication conflict.

We measured the distribution of the number of pauses per gene in the highest quartile of transcription and observed a rapidly decreasing curve (Fig. 3B). This indicates that the pause enrichment is not arising from a minority of genes that are highly problematic compared to a random genomic locus, but from a more uniform increase in pausing around many genes in the most-transcribed set. We chose a gene at random from the first quartile of transcription, and visualized one molecule with a co-directional replication track registering a pause close to its transcription start site (Fig. 3C). This was rescued by an incoming replisome passing over the gene. For comparison, we show a co-directional track with no pauses over this locus; it terminated normally with an incoming fork (Fig. 3D).

Finally, to confirm that ± 250 bp is a reasonable length scale to probe enrichment, we determined the spatial variation in enrichment across the start/end sites at higher spatial resolution (see *Methods*). In either relative direction, enrichment peaks within ∼100 bp at either end of a gene and is somewhat depleted over the gene body (Figs. 3E, S6C). The peaks in pause enrichment are just upstream of the TSS consistent with a promoter-proximal impediment to replication. By contrast, pause enrichment proximal to the TTS is predominantly within the transcribed region, consistent with RNAPII-replication conflicts. Thus, we see that RNAPII-transcribed elements pose a threat to replisome progression in a transcription-dependent manner and that head-on conflicts are worse than co-directional, particularly at the transcription end site.

### R-loop level is a stronger predictor of replisome pausing than RNAPII occupancy

We further investigated our observation that replisome pauses were more enriched at termination sites rather than start sites of RNAPII-transcribed genes. A previous study in budding yeast reported that R-loops are also more enriched at termination sites on average^39^. Thus, we repeated our earlier analysis but using R-loop level instead of RNAPII occupancy to place genes in quartiles. We used population-level RNase H1 CRAC measurements on asynchronous cells from ref.^38^, which was shown to be a proxy for R-loops. The set of pauses and genes are almost identical from the previous subsection, but due to the re-binning based on R-loop level, a minority of genes get shuffled into different quartiles (Figs. S7A, B). The mean spatial profiles of RNAPII and R-loop level across genes are consistent with previous measurements (Fig. S7C)^38,39^.

At start sites, we saw that pause enrichment of both relative directions were approximately equal overall and correlated with R-loop level on average, similar to the analysis above for RNAPII occupancy (Figs. 4A, S8A, B). However, at termination sites, head-on pause enrichment correlated better with the level of R-loop signal than the level of RNAPII occupancy. Enrichment was more pronounced in the first R-loop quartile as opposed to RNAPII (∼1.4x vs ∼1.35x), and then sharply dropped and plateaued with decreasing R-loop signal, unlike the more uniform decrease for RNAPII (Figs. 3A, 4A).

**Figure 4.**
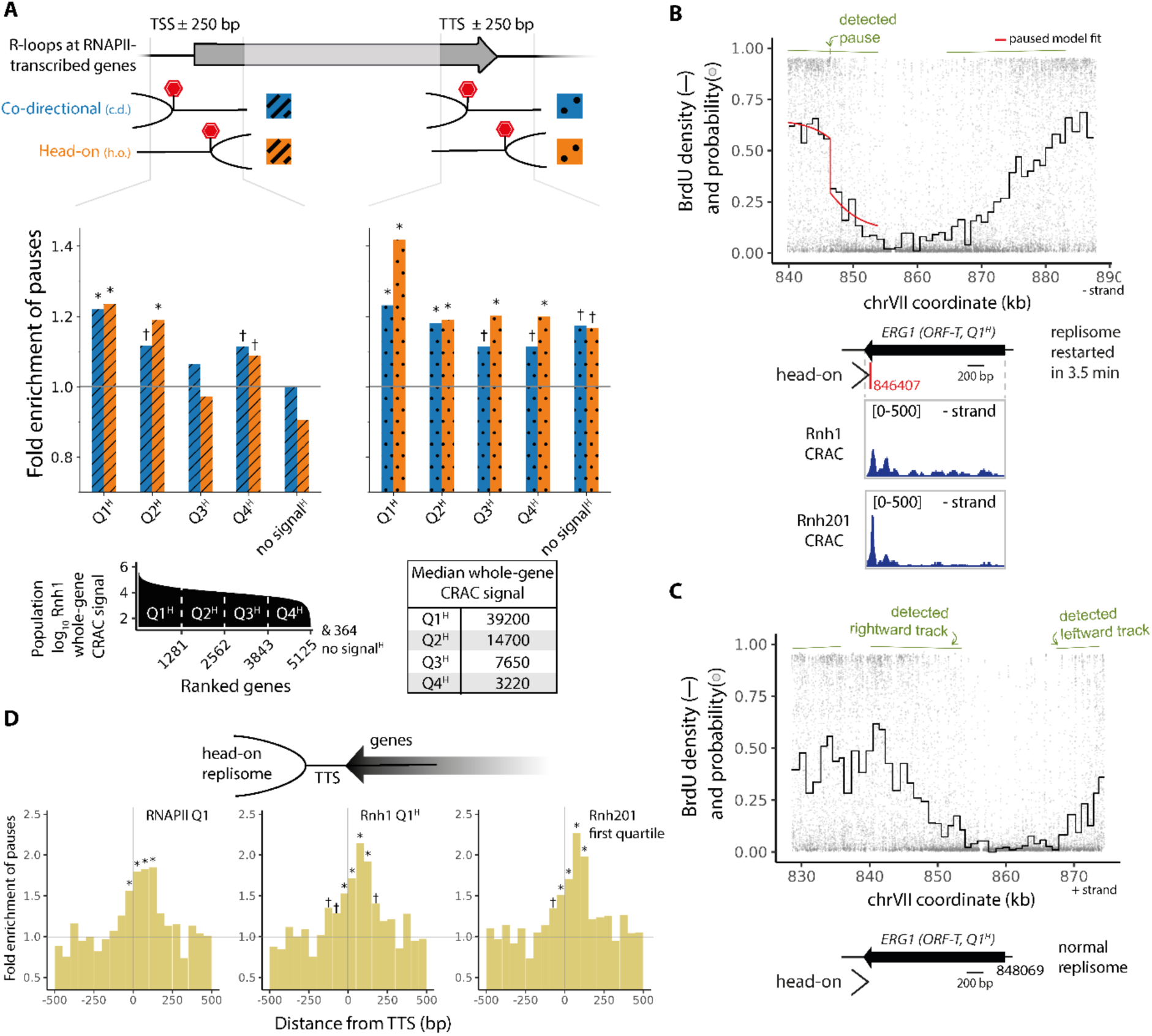
Replisome pause enrichment at termination sites of RNAPII-transcribed genes correlates better with Rnh1 than RNAPII occupancy. **(A)** Top: Ratio of observed to expected pause count versus quartiles of total amount of Rnh1 occupancy per RNAPII-transcribed gene at transcription start and termination sites (TSS and TTS respectively). Pauses counted in 500 bp regions centred at TSS/TTS. Pauses classified as head-on or co-directional (see *Methods* and Fig. S8A for *n* values of pauses). Bottom left: Whole-gene Rnh1 occupancy signal versus genes ranked in descending order. Bottom right: Table of median whole-gene Rnh1 occupancy signals per quartile. **(B, C)** Two example single-molecule BrdU density profiles of molecules displaying head-on replisome tracks over the *ERG1* gene, one with a pause within the gene ∼70 bp from its transcription end site that restarts in 3.5 min **(B)** and one without any pauses **(C)**. Red: best-fit model with pause. Green: replisome track annotations. Bottom of **(B)**: line diagram of pause location and replisome orientation relative to gene and nature of replication rescue, and Rnh1, Rnh201 occupancy signals over the gene. **(D)** Ratio of observed to expected pause count versus relative distance of head-on pauses from nearest TTS in 50 bp bins. Only pauses near genes in highest quartile of total RNAPII, Rnh1, or Rnh201 occupancy are included (left to right, *n* = 852, 869, and 895 pauses respectively). Throughout, * and † indicate > 3 s.d. or > 1.65 s.d. respectively of enrichment. Genomic coordinates from W303 reference^59^.

We chose a gene at random from the first quartile of R-loop signal and visualized one molecule with a head-on replication track registering a pause close to its transcription end site (Fig. 4B). The synthesis restarted in ∼4 min, proceeded for another ∼7 kb or equivalently 3.5 min, before terminating with an incoming replisome. The pause site coincided with a peak in R-loop CRAC density. We show a head-on track with no pauses over this locus as well, which appeared to terminate normally with an incoming fork (Fig. 4C).

For head-on pauses, the spatial enrichment over the termination sites of the top quartile genes showed higher peaks in the R-loop analyses than in the RNAPII analysis – this was apparent for both RNase H1 and RNase H2 CRAC signal (Figs. 4D, S8C). In summary, our data are consistent with R-loop level being a better predictor of head-on replisome pausing than RNAPII occupancy at transcription termination sites.

### Replisome pause enrichment at SPT5 proximal to a G4-forming sequence

Although most replisome pause events are infrequent and distributed across the genome a minority are enriched at certain genomic loci. Using our measurements of expected and observed pause counts over the genome, we identified zones of significant enrichment (515 zones, enrichment ≥ 3 s.d., Table S4). Below we highlight two example loci with significant enrichment in replisome pauses.

The highest peak in pause count on chromosome XIII occurred over the transcription termination site of the *SPT5* gene (Figs. 5A, B). This gene is in the top quartile of both RNAPII and R-loop occupancy, but its termination site was a clear outlier in pause count among the highly transcribed genes (Fig. 3B). This suggests a process other than transcription may contribute to pausing here. Over a 7 kb region, centred on the *SPT5* gene, we detected 689 replisome tracks, of which 49 show pauses within this region (∼7%; Fig. 5B). All 49 pauses were on leftward replisomes and were largely distributed over two zones of pause enrichment ∼2-3 kb long, one overlapping the *SPT5* transcription start site (zone 1) and the other the termination site (zone 2; Fig. 5B).

**Figure 5.**
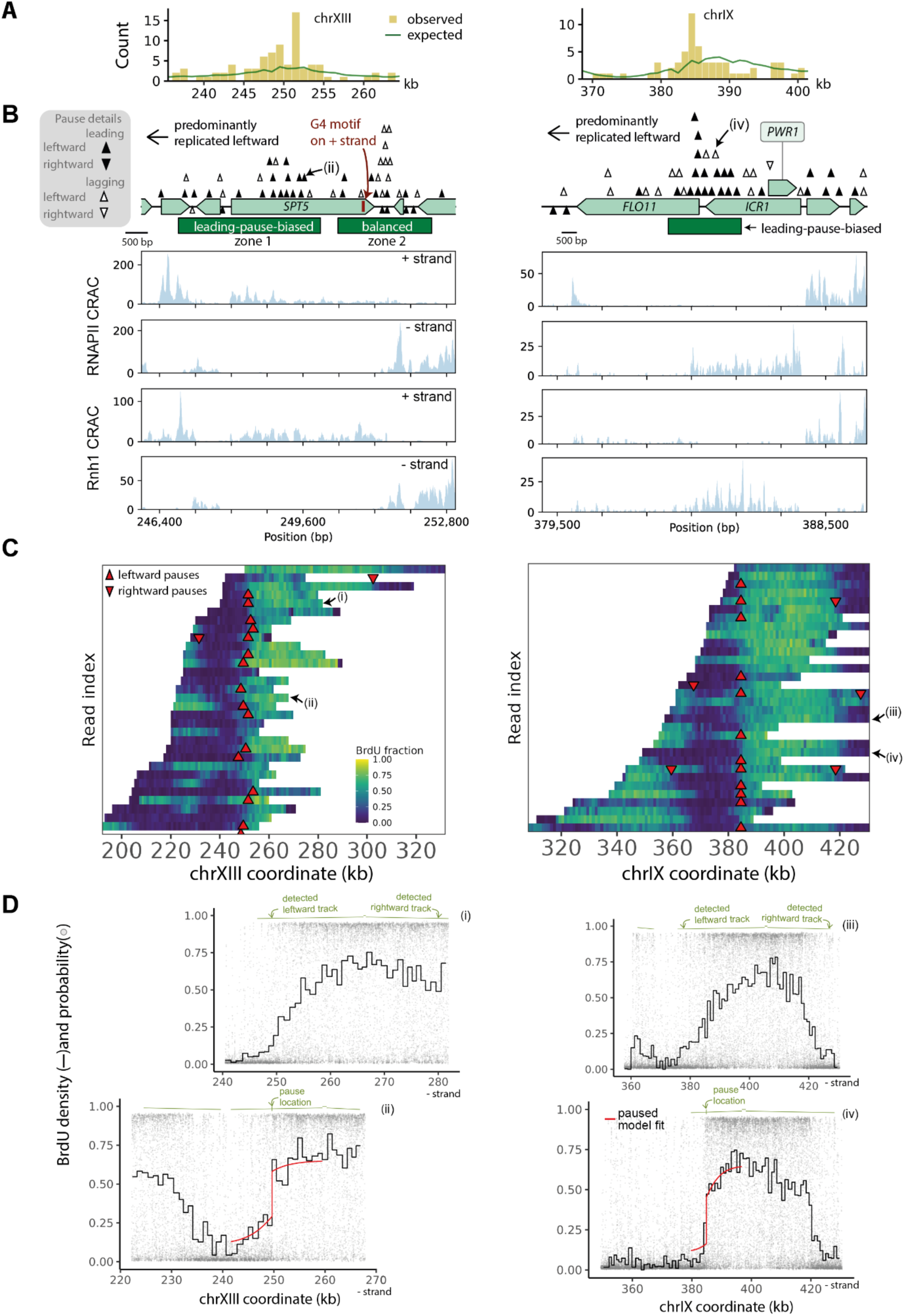
A bias towards pausing of leading strands is consistent with helicase-polymerase uncoupling at a G4 motif and a long, non-coding RNAs. **(A)** Expected and observed count of single-molecule replisome pauses versus mapped location along sections of chromosomes XIII and IX. **(B)** Top: locations of pauses (arrowheads) detected at *SPT5* and *FLO11* loci, with annotations of zones with > 3 s.d. pause enrichment (dark green rectangles) and transcribed regions (light green). Middle and bottom: RNAPII and Rnh1 occupancy signals at each locus. Arrowheads coloured black or white indicate leading or lagging-strand pauses, respectively, and up or down direction indicates leftward or rightward replisome pauses, respectively. Highlighted molecules (ii) and (iv) over *SPT5* and *ICR1* are visualized in **(C)** and **(D)**. **(C)** Heatmaps of BrdU density for a subset of molecules with and without pauses overlapping *SPT5* and *FLO11*. Non-paused molecules were selected at random, and paused molecules were either selected at random from the locus of chrXIII:246000-253000 or represent the entire set from chrIX:384000-385000. **(D)** Single-molecule BrdU density versus genomic coordinate of four molecules overlapping with *SPT5* or *FLO11*. Two are unperturbed and two contain pauses—4.0 min pause head-on relative to *SPT5* transcription followed by rescue from incoming replisome (left), and a 4.2 min pause co-directional relative to *ICR1* transcription (right). Genomic coordinates from W303 reference^59^.

Interestingly, a previous study highlighted the *SPT5* gene as containing an exemplary G-quadruplex (G4) motif near its termination site^3^. The site displayed high occupancy of the Pif1 helicase and paused replication, as seen with a population measurement, when the nuclear form of Pif1 was absent. For ease of discussion, whenever we refer to a G4 in this section, we are referring to this short sequence which has a propensity to form a G4 as also reported in another study^40^. The G4 is present on the top strand close to the left-end of our zone 2 (Fig. 5B). As this entire region is predominantly (90%) replicated leftward, the G4 lies on the leading strand template. Interestingly, leading and lagging strand syntheses showed different pause likelihoods on either side of the G4. A leftward replisome would replicate most of zone 2 before arriving at the G4; the pauses here are clustered ∼500 bp ahead of the G4 and were balanced between leading and lagging strands (10 and 12 respectively). As synthesis moves ∼1-2 kb beyond from the G4 (zone 1), the pauses were predominantly from leading strands—17 as opposed to 7 from lagging strand. Such biases and higher only occur with a ∼3% rate from random chance (see *Methods*).

We visualized a subset of tracks with and without pauses over this locus using a heatmap and single-molecule views (Figs. 5C, D). The paused event in Fig. 5D was chosen from the leading-strand cluster over zone 1 (Fig. 5B); this leftward synthesis paused for 4.0 min before restarting and continuing for another ∼8 kb (equivalently to 4 min) before terminating with an incoming rightward replisome. In summary, we observe enrichment of pauses along the path of synthesis behind and ahead of a previously annotated G4 site in a highly transcribed gene, with the pauses after the G4 encounter predominantly being observed on leading strand synthesis.

### Replisome pause enrichment near *FLO11* correlates with a long, non-coding RNA

The highest peak in pause count on chromosome IX occurred near the transcription start site of the *FLO11* gene (Figs. 5A, B). The pauses could be due to *FLO11* transcription, but this is reported to be low in W303^41,42^; consistently, we measured that *FLO11* belongs to the lowest quartile of transcription and R-loop signal. However, its ∼3 kb promoter is transcriptionally active^43^ with two long, non-coding RNA genes—the ∼3 kb *ICR1* and ∼1 kb *PWR1*. These form an epigenetic *FLO11* toggle—the co-directional *ICR1* transcription represses it, whereas the oppositely-oriented *PWR1* activates it^44^. Our observations are consistent with epigenetic suppression of *FLO11* expression and indicate a prominence of R-loops across a ∼3 kb regions corresponding to *ICR1* (Fig. 5B).

Consistent with replisome pausing due to *ICR1*, we see a ∼2.5 kb pause enrichment zone near *FLO11* that intersects with the high RNAPII and R-loop occupancy associated with *ICR1* (Fig. 5B). Most replisomes are unimpeded—586 tracks pass through the locus (Fig. 5B), of which 43 contain pauses (7.3%) and 21 are present within the enrichment zone (3.6%). Thus, we are detecting rare, spatially diffuse pauses for which our technique is well-suited. Almost all pauses are co-directional with *ICR1* transcription since this region is primarily (∼90%) replicated leftward. We visualized a selection of paused and non-paused tracks in a heatmap, from which we display two example single molecule views: one without pausing, and one where synthesis paused for 4.2 min (Figs. 5C, D). How synthesis was resumed on this molecule is unclear—either fork restart or rescue from an incoming fork are consistent with the data.

Within the pause enrichment zone, we observe a strong bias towards pausing on leading rather than lagging strands (Fig. 5B, 16 out of 21 pauses). Such a bias or higher occurs only at a ∼1% rate from random chance (see *Methods*). The top strand is the template for *ICR1* transcription, is bound by resulting RNA in R-loops, and as replication is predominantly leftward here, the top strand is also the template for leading-strand synthesis. Therefore, this could explain the leading-strand bias in pauses. In summary, we have identified replication pauses over the promoter of the epigenetically controlled *FLO11*.

## Discussion

Here we present a quantitative map of replisome pauses across the budding yeast genome. On single sequenced molecules, local BrdU density served as a measure of replication time and sharp drop-offs identified replisome pauses. For each single molecule replisome pause, from the spatial BrdU density profile, we can determine the direction of replisome movement, the leading/lagging-strand identity, the location and approximate duration of the pause and whether the replisome resumed synthesis. This has led to new insights. For instance, in the well-studied unidirectional programmed replisome barrier at the rDNA, we measured a ∼120 bp separation between the average leading and lagging-strand pause site, which is similar to the size of an Okazaki fragment (Fig. 1E).

Globally, we saw a surprisingly high pause rate— ∼5% of detected replisomes in the non-repetitive genome pause, with a majority of pauses diffusely distributed across the genome (Figs. 2A, B, S4). This implies that the yeast cell faces up to ∼13 pauses in a single S phase (based on ∼250 concurrent replisomes^45^), which are spatially widely distributed. Single-molecule techniques are well-suited to capture such spatially diffuse and infrequent events. They complement population averages traditionally used in studies querying replication problems^3–7,12–14,19,21^. Such studies enrich for loci with highly localized events by design, and then may conflate short, frequent pauses with long, rare pauses at each locus as has previously been discussed^19^.

Although replisome pauses are largely diffuse over the whole genome, they are significantly enriched over transcribed features such as tRNAs, protein coding genes, and the 35S gene within the rDNA (Figs. 1-3). We determined that head-on conflicts were more numerous than co-directional using tracks with single-molecule replisome direction. We measured the following enrichment of head-on conflicts compared to a random genomic locus: ∼4x at the 35S rDNA gene in *fob1Δ* cells, ∼2x at tRNA genes, and ∼1.4x at the transcription termination site of highly transcribed protein coding genes. It must be noted that these are markedly less pronounced than the programmed fork barrier at the rDNA in wildtype cells, where rightward replisomes paused at ∼50x the level expected from a uniform distribution across the rDNA repeat.

Why are replisome pauses enriched at transcribed elements? Our results suggest that multiple phases of transcription may contribute. Firstly, a partially or fully assembled and non-transcribing complex may pause replisomes, as we observe pause enrichment upstream of transcription start sites (Fig. 3E) and at a nuclear peripherally localized ETC site that binds the transcription factor TFIIIC but does not initiate transcription (data not shown)^46^. Similarly, a previous study showed TFIIIB hinders fork progression even without Pol III transcription in yeast lacking the DNA helicase Rrm3^47^. Secondly, either transcription itself or an ensuing R-loop or both may drive conflicts with replication. We observe pause enrichment at start and termination sites of RNAPII-transcribed genes, and the pause rate increased for higher levels of transcription (Fig. 3A). Head-on conflicts at termination sites were the most problematic (Figs. 3A, B, E). Interestingly, this terminal pause enrichment correlated more strongly with R-loop abundance than with RNAPII (Fig. 4). Two possibilities are – (1) R-loops, perhaps with associated transcription complexes, are a greater challenge to replisomes or (2) R-loops form due to conflict or are stabilized or selected by it; i.e. R-loops may be a cause or a consequence of replication stress as previously reported^48–50^ and reviewed^51^.

The strand of the sequenced nascent DNA molecule and the direction of replication allow us to distinguish between leading and lagging strand synthesis. At the rDNA we observe similar numbers of leading and lagging strand pauses consistent with pausing of the whole replisome. Likewise, across the transcription termination site of the highly expressed *SPT5* gene we observe an increased frequency of replication pausing for both leading and lagging strands (Fig. 5B). However, within the *SPT5* gene we observe a strong bias towards leading (over lagging) strand pauses (Fig. 5B). This pattern indicates pausing of the leading strand polymerase, epsilon, while lagging strand synthesis continues, consistent with uncoupling of the helicase from polymerase epsilon. Therefore, using ensembles of the detected single-molecule replication pauses we can see a signature of helicase-polymerase uncoupling at rates as low as ∼1 in 20 replisomes.

The pronounced leading strand pause site within the *SPT5* gene is proximal to a strong G-quadruplex (G4) forming sequence that has previously been reported to slow replication when the nuclear form of the Pif1 DNA helicase is depleted^3^. This prior study indicated that replication was perturbed both in approach and beyond from the G4 motif. Consistent with this, we observe replication pauses on both sides of the G4 within *SPT5*, but upon movement beyond the G4 there is a strong bias towards leading strand pauses. This is consistent with the pausing on approach to the G4 affecting the whole replisome, while upon passage through the G4 there arises a risk of helicase-polymerase uncoupling. It is notable that the leading strand pauses span a region located 1-2 kb beyond the G4. This may indicate that leading strand synthesis can continue a short distance after helicase uncoupling before DNA synthesis stops (Fig. 6).

**Figure 6.**
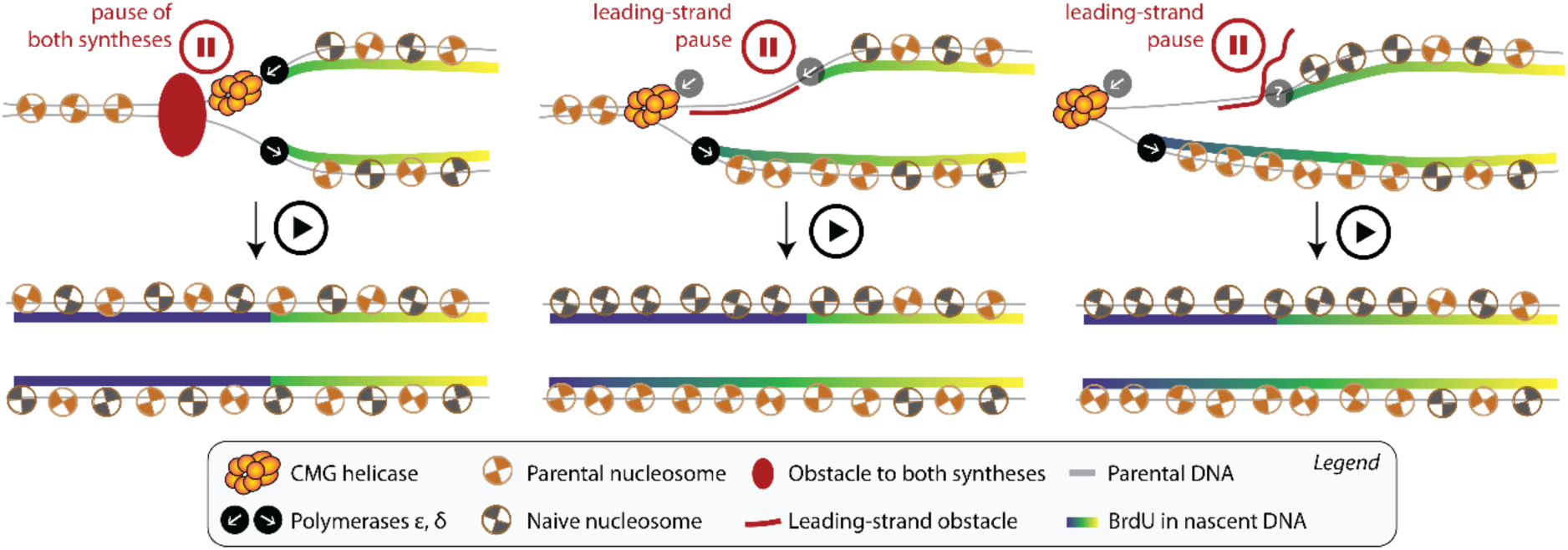
A model for distinguishing replisome from leading strand polymerase obstacles. Left: obstacles that block the whole replisome, for example blocking DNA unwinding by the CMG helicase, result in pauses to both leading and lagging strand synthesis. Our data at the rDNA replication fork barrier indicate that leading strand synthesis can continue to ∼35 nts of the barrier, whereas, lagging strand synthesis pauses ∼150 nts behind the barrier. Centre: by comparison, strand specific obstacles, e.g. R-loops or G4s, could block polymerase synthesis while allowing other replisome components to proceed. Blocks to lagging strand synthesis would be readily bypassed due to the continuous repriming of Okazaki fragments. However, blocks to leading strand synthesis could uncouple the leading strand polymerase (ε) from the helicase. In this scenario lagging strand synthesis could continue while leading strand synthesis is paused. Consequently, parental histones (with associated post-translational modification) would only be inherited by the lagging strand. Once replication is completed the leading strand would be populated by naïve histones lacking appropriate epigenetic marks. Right: leading strand synthesis may be able to continue a short distance after the uncoupling event before stopping. This mechanism could explain the displacement of detected replication pause sites from presumed leading-strand obstacles.

Replication pauses and the uncoupling of helicase and leading strand polymerase have been suggested as the molecular event that triggers epigenetic switching in eukaryotes^52^. However, to our knowledge, replication pausing has never been observed in the vicinity of an epigenetically regulated gene. Here we report pausing in replication of the leading strand upstream of the epigenetically regulated *FLO11* gene. This region contains a long non-coding RNA (*ICR1*) and R-loop, associated with the leading strand template, that may be responsible for helicase-polymerase uncoupling (Fig. 5B). Consistent with helicase-polymerase uncoupling are ChIP-chip data indicating enrichment of polymerase epsilon subunits in this region^7^, but not GINS subunits, which were otherwise enriched at synthesis-stalling hotspots such as tRNA genes^19^. Helicase-polymerase uncoupling is proposed to disrupt the redistribution of nucleosomes from ahead of the replication fork to the newly synthesised leading strand (Fig. 6). After delay, completion of leading strand synthesis would use packaging with naïve nucleosomes; thus, destabilizing epigenetic inheritance of one daughter DNA molecule. This is consistent with genetic evidence from budding yeast^53^, Arabidopsis^54^, and cultured chicken cells^55^. Therefore, we have, for the first time, observed the potential molecular trigger for eukaryotic epigenetic switching – replication perturbation and helicase-polymerase uncoupling – upstream of an epigenetically regulated gene.

In summary, we have presented a single-molecule modality for replisome pause measurement. Our approach allows a highly quantitative and sensitive determination of where and for how long DNA synthesis is paused. This allowed us to identify rare, but important, pauses in DNA replication that have been missed by cell population-based techniques. This includes and distinguishes between pausing of the whole replisome and pausing of one strand of synthesis. Recent advances in the application of single-molecule mapping of replication dynamics in a range of organisms^18,23,56,57^, including human cells^58^, highlights the potential of our approach. In the future, increases in sequencing depth and improvements in base analogue detection will further the identification of rare replication events that have a major impact on cellular and organismal phenotypes.

## Acknowledgement

We thank Julian Blow, Anthony Carr, Jamie Carrington, Isabel Díez Santos, Anne Donaldson, Peter Gillespie, Shin-Ichiro Hiraga, and Alexandra Pyatnitskaya for helpful discussions and comments on the manuscript. This work was supported by the Biotechnology and Biological Sciences Research Council (BBSRC), part of UK Research and Innovation, through the Core Capability Grant BB/CCG2220/1 at the Earlham Institute and the Earlham Institute Strategic Programme Grant Cellular Genomics BBX011070/1 and its constituent work packages BBS/E/ER/230001B (CellGen WP2 Consequences of somatic genome variation on traits). The work was also supported by the following response-mode project grants: BB/W006014/1 (Single molecule detection of DNA replication errors) and BB/Y00549X/1 (Single molecule analysis of Human DNA replication). This research was supported in part by NBI Research Computing through use of the High-Performance Computing system and Isilon storage. Part of this work was delivered via Transformative Genomics, the BBSRC funded National Bioscience Research Infrastructure (BBS/E/23NB0006) at Earlham Institute, by members of the Single-Cell and Spatial Analysis Group.

## Author contributions

CAN conceived the study. CAN and ST designed the study. AR and CAM designed experiments with CAN and performed them. ST, CAN, and AR wrote scripts, analysed the data, and prepared figures. ST and CAN wrote the draft with inputs from all authors.

## Data and code availability

Data is archived in Zenodo under the DOI 10.5281/zenodo.16784880. Code is available at https://github.com/DNAReplicationLab/fork_arrest. Scripts were written in shell script, Python, and R.

### Video Legends

**Video 1**. Schematic of sister replisomes moving leftward and rightward from their initiation site over time in a cell incorporating BrdU into nascent DNA. When BrdU availability relative to thymidine decreases over time, the cell records fork movement as spatial gradients in BrdU density along the newly synthesized DNA.

**Video 2.** Schematic of sister replisomes as in Video 1 but now the leftward-moving fork temporarily arrests at a site. The delay due to arrest results in a lower amount of BrdU available to the fork at restart, thus creating a sharp spatial decline in BrdU density along the DNA at the arrest site against a smoother background decline.

## Methods

### Strain construction

Yeast strains used in this study are from the W303 background and described in Table S1. Yeast transformations were performed following the lithium acetate method^60^. The *fob1Δ* strain was made by PCR amplification of the KanMX knockout cassette from the Yeast Deletion Collection and integration into the genome^61^. Strain construction of ARY017 was performed using a custom CRISPR-Cas9 plasmid (pAR021) and healing fragment made from NotI digested pAR045 containing budding yeast codon optimized versions of *hENT1* and *hsvTK*. pAR021 and pAR045 were made using parts from Yeast MoClo kits and custom plasmids from GenScript^62,63^.

### Yeast growth and cell cycle experiments

Cell cycle experiments and DNA extraction for the *fob1Δ* strain were performed as previously described^23^. Strain ARY017 was grown in YPAD (Formedium, CCM1010) at 30°C to early log phase (OD600∼0.300) and arrested in G1 with alpha factor (final concentration 0.5 µM, GenScript, RP01002). BrdU (Sigma-Aldrich/Merck, B5002) was added to a final concentration of 30 µM 25 minutes prior to S-phase release with pronase (final concentration 200 µg/ml, Millipore/Merck, 53702). 20 minutes into pronase treatment, nocodazole (final concentration 15 µg/ml, Sigma-Aldrich/Merck, 487928) was added to arrest cells in G2/M preventing entry into another cell cycle. Cells were collected 90 minutes after pronase addition for DNA extraction. HMW DNA extractions were performed using a modified PacBio Nanobind Tissue Kit protocol (PacBio, 102-302-100).

### Flow cytometry

Standard protocols were followed for preparation of cells in SYTOX Green for flow cytometry^64^. Briefly, cells were washed and stored in 70% EtOH during the cell cycle experiments. Cells were washed twice in 50 mM Sodium Citrate, resuspended in 1 ml 50 mM Sodium Citrate and RNase A (final concentration 0.25 mg/ml, MP Biomedicals, 219398050) and incubated at 50°C for one hour, 20 µl of 20 mg/ml Proteinase K (MP Biomedicals, 193504) was added and again incubated at 50°C for one hour, finally cells were resuspended in 1 ml 50 mM Sodium Citrate with 1x SYTOX Green (Invitrogen, S7020). Immediately before flow cytometry, cells were briefly sonicated and strained (Falcon, 352235). 50,000 cells were collected on the BD FACSAria Fusion with 488nm laser excitation with 530/30 emission filter. Flow cytometry results were analysed using FlowJo v10 (BD Life Sciences).

### Nanopore sequencing

DNA was prepared for sequencing following protocols from Oxford Nanopore Technologies (ONT). Ligation library preparations were performed for all samples (ONT, SQK-LSK109). The nanopore library for the *fob1Δ* strain was sequenced on a MinION Mk1 sequencer using flow cell version R9.4.1 as previously described^23^. Three libraries were prepared for ARY017 and sequenced on PromethION 2 Solo (ONT, PRO-SEQ002) using one PromethION Flow Cell R9.4.1 (ONT, FLO-PRO002). Each library was run for approximately 24 hours, then the flow cell was washed (ONT, EXP-WSH004), and a fresh library was loaded.

### Measurement of BrdU substitution and detection of replication tracks

We started from two nanopore datasets from sequencing of wildtype cells—one from our previous study^23^ (labelled ‘wildtype_independent’), and a larger dataset generated here (unlabelled or labelled ‘wildtype’). We also used data from *fob1Δ* cells wherever indicated. We obtained DNA sequences from the recorded currents using the ONT basecaller guppy with the config file *dna_r9.4.1_450bps_hac.cfg*. We then aligned sequences to a reference genome using minimap2^65^ (see the next paragraph for reference genome) and performed reference-anchored modification calling per thymidine per molecule using DNAscent v2^26^. We converted DNAscent outputs to the modBAM format with DNAscentTools (https://github.com/DNAReplicationLab/DNAscentTools), and retained only molecules with an alignment length greater than or equal to 30 kb for further analysis. We used a threshold substitution probability of 0.5 to call individual thymidines as BrdUs in calculations.

For alignment and modification calling in the non-repetitive genome, we used a W303 reference from a previous study after removing the rDNA contigs, the mitochondrial chromosome, and the 2 micron plasmid, and renaming chromosomes to the conventional roman numeral style used in *sacCer3*^59^. For the rDNA results, we made and used a custom reference with just the rDNA repeat sequence (chrXII:451686-460822 from *sacCer3*, both coordinates inclusive) repeated 22 times.

In the non-repetitive genome, we detected replication tracks on single-molecules and assigned direction (left/right) using DNAscent forkSense v2, whereas in the rDNA we first detected replisome pauses and assigned the direction of the BrdU gradient at the pause site to the direction of the paused replisome. For some analyses, we further identified replisome tracks as those belonging to leading (right, + or left, -) or lagging strand synthesis (left, + or right, -) processes where the two entries in parentheses are the replisome direction and the strand alignment respectively.

### Relation between BrdU density and replication time

We lifted over ensemble measurements of S phase DNA copy numbers, which were measured using Sort-seq, and median replication time from ref.^66^ using liftOver^67^. These data were in 1 kb genomic windows, so we measured mean BrdU density per window per molecule passing through each window and analysed the relation. We measured a negative correlation between median replication time and mean BrdU density, and a positive correlation between mean BrdU densities and copy numbers (Fig. S1E-G). The two relations are opposite to each other as median replication time decreases with copy numbers^66^. Both analyses are consistent with previous studies that used similar strains^23,24^, and show that BrdU incorporation in nascent DNA decreases with time from S-phase onset.

#### Single-*molecule* replisome pause detection

In molecules aligned with the rDNA, we selected those with a mean BrdU density of at least 0.05 and measured the density difference between an upstream and downstream window at each thymidine per molecule (window size 3x 290T or approximately 3 kb in the yeast genome). Then, we detected peaks in the density difference, requiring that peaks are at least 15x 290T apart on the genome (approximately 15 kb). Then, we collated peaks in density difference across all molecules, filtering out those with absolute values of the density difference smaller than 3x the s.d. of all density differences across all molecules. We also filtered out those peaks which were closer than 5 kb to the ends of molecules, and those with maximum and minimum BrdU densities ≤ 0.5 or ≤ 0.01 respectively in their upstream or downstream windows. Thus, we obtained our rDNA pause dataset. We observe a similar pause pattern irrespective of whether we used a 3 s.d. or a 2 s.d. threshold (Figs. 1E, S2A). We calculated the modal leading and lagging strand pause site as the midpoint of the bin with the most pauses in Fig. 1E.

In molecules aligned with the non-repetitive genome, we obtained a best-fit model first to establish typical BrdU patterns across a normal replisome track, so that we can use it as a reference to detect pauses. We assumed BrdU fall-off across a normal replication track followed a sigmoidal shape based on our visualization of a few tracks (see eq. (1) below for the functional form of the sigmoid). We used forkSense-called replisome tracks with min length 10 kb on molecules with minimum alignment length 30 kb. We performed model fitting in batches—by sampling 1000 tracks at a time, windowing each track using non-overlapping windows of 300T each, rejecting tracks without a gradient (with an auto-correlation length of less than 2 windows or equivalently ∼2 kb), and searching from a grid of low, high, width combinations which sigmoid best-fit all the tracks collectively (Figs. S3A, D, E). We then averaged parameters from many fits to get the parameters of the reference sigmoid. The functional form of the sigmoid along a detected replisome track is

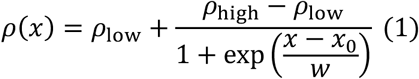

where ρ is the BrdU density, *x* is the genomic coordinate along the replisome track, and ρ_low_, ρ_high_, *w* and *x*_’_ are the low, high, width, and the centre of the sigmoid respectively. Right- and left-moving tracks have positive and negative *w* respectively. The centre *x*_0_ is specific to each track. We see a consistent gradient across replication tracks in our data here and also from analysis of molecules from our previous study (Figs. S3A, D, E).

After obtaining a reference sigmoidal model, we proceeded to detect replisome pauses using a ‘cut-and-align’ approach i.e. we ‘cut’ a detected replisome track at every thymidine and fit two copies of the reference sigmoid obtained above separately to the thresholded BrdU calls in the upstream and downstream sections (‘align’). We used a standard generalized linear model (GLM) fitting approach implemented in Tensorflow with only the centres *x*_’_ of the two sigmoids as fitting parameters. We then identified one pause site per track as that thymidine at which the two sigmoids put together achieved the best fit, and the associated pause duration as the distance between the corresponding two best-fit centres (Fig. S3B). We performed this only on tracks longer than 3 kb on molecules with alignments longer than 30 kb, discarding those tracks where errors in fitting occurred at or above 3% of thymidines. We then further filtered the list of pause sites to avoid false positives; the primary filters are a positive pause duration and a high confidence on the location of the best-fit pause site (Supplementary Note 2). We verified the method by comparing measured and actual pause sites and durations on simulated replisome tracks (Supplementary Note 2).

We then calculated the expected pause count versus genomic coordinate, which is our null hypothesis or, equivalently, the pause sensitivity. We gathered replisome tracks with pauses, and the upstream and downstream reference sigmoids fit to them (see paragraph above) and calculated the statistics of the location of the pause site relative to the centre of the reference sigmoid upstream of the pause site. We then gathered the centres of the sigmoids fit to replisome tracks with no pauses, placed them on the reference genome, and superimposed the distribution calculated in the previous step relative to each centre along the reference genome. In other words, we use our detected pause locations to simulate where they could have been found on normal forks. We added all the superimposed functions, and normalized to total pause number, thus producing a landscape of pause sensitivity. We repeated this calculation separately for left and right tracks so that we can calculate head-on and co-directional pause enrichment with respect to genes.

### Visualizations

To plot heatmaps, we measured BrdU density along individual molecules in regular 1 kb windows along the reference genome. Then, we stacked horizontally and vertically a randomly selected subset of those molecules with a mean BrdU density greater than 0.05. To aid in visualization, our algorithm chose a stacking order for the molecules such that neighbours on the same row were at least 5 kb apart, and this appears as white bands. For single-molecule figures such as Fig. 1D, we plotted raw (grey), and averaged (black) modification calls, and optionally superimposed model fits at pause sites (red) and forkSense-called features (green). To calculate the average BrdU density across a molecule, we used non-overlapping windows with 300 thymidines each, with boundaries registered to the left end of the molecule. We plotted pause counts presented as a fold enrichment in 13 bp bins in Figs. 1E, F, S2A, F (wildtype) and in 60 bp bins in Figs. 1H, I (*fob1Δ*); a smaller bin size was needed for the wildtype pauses as they were highly clustered at the rRFB. To calculate fold enrichment in the rDNA, pause counts per bin were divided by expected counts from a spatially uniform distribution; this was done separately for each kind of pause in the figure (left/right or lead/lag). To plot genomic datasets, annotations, and measured pauses stacked vertically (Fig. 5B), we used a modified version (https://github.com/sathish-t/DnaFeaturesViewer) of the DNAFeaturesViewer package^68^. We visualized annotations from SGD in Fig. 5B excluding those described as proteins of unknown function or dubious open reading frames. We visualized CRAC profiles in Fig. 3C, 4B using IGV^69^. To visualize CRAC data, we first used bigWigToBedGraph^70^ to convert data to plain text and then lifted over to W303 coordinates. We used RIdeoGram to plot expected and observed pause counts across the genome^71^ (Fig. S4) and deepTools to plot CRAC signals versus scaled distance along transcripts on *sacCer3* coordinates^72^ (Fig. S7C).

### Genomic annotations used for correlation with pause sites and calculation of enrichment

We obtained coordinates of the replication origin and the 35S gene in the rDNA from SGD^73^, and the Fob1 binding sites from ref.^27^. In the non-repetitive genome, we obtained tRNA locations using tRNAscan-SE on our reference genome^74^. We ranked RNAPII-transcribed genes into quartiles by summing the raw Rpb1 (a subunit of RNA polymerase II) or Rnh1 (RNase H1) CRAC (crosslinking analysis of cDNAs) signal from ref.^38^ (dataset labels S45_R2 and Asyn_R2 respectively) with strand-specificity over transcript coordinates, removing overlaps with tRNAs and the rDNA region if needed. We used ORF-transcript coordinates measured in ref.^15^ and stored in SGD. We also used Rnh201 signal (subunit of RNase H2, dataset label Asyn_R2) for one panel in Figs. 5B and S7C.

To calculate spatial enrichment of pauses around a set of features e.g. transcription end sites of genes, we lifted over the features to our reference genome. We measured the signed distance between every base on both strands and the nearest feature and segmented the genome into bins of this relative distance. We then split the bins into quartiles according to some signal if needed e.g. transcription level of the nearest gene, and calculated expected and observed pause counts, separately by the relative direction between the underlying replisome tracks and the nearest feature. We used the standard packages of samtools^75^ and bedtools^76^ to perform this counting. We formed zero-signal bins of Figs. 3, 4 using the few transcripts with no signal and the set of no-signal genes identified by summation of CRAC signal but now using the coding sequence coordinates in SGD instead of transcript coordinates.

### Statistical analyses

To evaluate statistical significance of pause enrichment over one or a set of genomic intervals, we added observed and expected pause counts over the interval(s) and calculated how many standard deviations the observed result is away from the expected result – i.e. the ratio (observed-expected)/(expected)^1/2^. The standard deviation of (expected)^1/2^ is a standard result from counting statistics. We then show if this ratio is above 3 s.d. (‘*’) or 1.65 s.d. (‘†’) in various plots; these correspond to a probability threshold of occurrence of 0.1% and 5% respectively by random chance from z-score tables. To calculate sites of pause enrichment as reported in Table S4, we added observed and expected pause counts in sliding windows of size 1 kb and stride length 10 bp over the non-repetitive genome, located windows with enrichment above 3 s.d. as described above, merged overlapping windows, and retained only those sites with at least two pauses. To determine the probability of *q* leading strand pauses and higher out of a set of *n* pauses in total, we used the formula 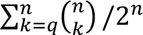 derived from the binomial distribution produced by *n* coin tosses.

