## Supplementary information for "Single-molecule landscape of DNA replication pausing"

* Correspondence should be addressed to Prof. Conrad A. Nieduszynski

**Table of contents**


- Supplementary notes 1 and 2
- Figures S1-9
- Tables S1-3
- Caption of Table S4 (table is a separate file)
- Supplementary references

**Supplementary note 1: Nearest-neighbour distances**

We measured the distribution of the distance between each detected pause and its nearest neighbour on the non-repetitive genome. We also visualized the expected distribution from the null hypothesis and from a uniform distribution of pauses over the genome, which we calculated as shown below.

*Expected nearest-neighbour distances*

Our input is the expected pause count $f(q)$ on non-overlapping, regular genomic windows whose positions are given by the coordinates $q$. This is our null hypothesis or a measure of the sensitivity of our technique. The probability density function $p$ that the closest neighbour of a pause at a genomic location $x$ is a distance $y$ away is given by

$$p(x,y)\Delta y = (f(x+M\Delta y)\Delta y + f(x-M\Delta y)\Delta y)\prod_{k=1}^{M-1}(1-f(x+k\Delta y)\Delta y)(1-f(x-k\Delta y)\Delta y)$$

Here, we discretize the reference genome using grid size $\Delta y$, and $M$ is the distance $y$ in terms of number of grid points i.e. $M = y/\Delta y$. We use the equation above to calculate the expected nearest-neighbour distance between pauses using the null hypothesis as $f(q)$ and averaging over all $x$ (Figs. 2B, S5B).

*Nearest-neighbour distances when pauses are uniformly distributed throughout the genome*

To leading order in $\Delta y$, $(1 - f(x+k\Delta y)\Delta y) = \exp(-f(x+k\Delta y)\Delta y)$. Using this simplification, we get

$$p(x,y)\Delta y = (f(x+M\Delta y)\Delta y + f(x-M\Delta y)\Delta y)\exp\left(-\sum_{k=1}^{M-1}(f(x+k\Delta y)\Delta y + f(x-k\Delta y)\Delta y)\right)$$

Taking the continuous limit of $\Delta y \rightarrow 0$, we get

$$p(x,y) = \big(f(x+y) + f(x-y)\big)\exp\left(-\int_{x-y}^{x+y} f(q)dq\right)$$

Thus, when the pause landscape is uniform i.e. $f(q) = N/L$ where $N$ is the total number of pauses and $L$ is the total length of the genome, the equation above simplifies to

$$p(y) = \left(\frac{1}{y^{\star}}\right)\exp\left(-\frac{y}{y^{\star}}\right) \quad (1.1)$$

where $y^{\star} = L/2N$ or half the mean distance between pauses. Thus, if all cells experienced identical problems uniformly throughout the genome, the probability $p$ of nearest-neighbour distances would follow equation (1.1) (Figs. 2B, S5B).

*Nearest-neighbour distances in our independent dataset*

We analysed spatial pause distributions in the molecules of our (smaller) independent BrdU dataset (1). Spatial patterns are noisier, with enrichment and depletion observed below and above ~1000 bp respectively, which is a length scale ~10-fold larger than the length scale of enrichment in our primary dataset (Figs. 2B, S5B). This can be explained from equation (1.1) above; the length scale of the uniform pause distribution $y^{\star}$ scales inversely with dataset size $N$. So, presumably our nearest-neighbour statistics also scale inversely with dataset size. In other words, as our independent dataset is ~8-fold smaller (Table S3), we expect Fig. S5B to resemble Fig. 2B if the X axis of Fig. 2B were stretched by a factor of 8, which is roughly the case. This suggests that we are observing comparable statistics in both datasets.

**Supplementary note 2: Filters used in the pause-detection procedure in the non-repetitive genome and pause detection in a simulated dataset**

After detecting one best-fit pause site per detected replisome track in the non-repetitive genome using our 'cut-and-align' approach (see *Single-molecule replisome pause detection* of *Methods*), we further applied filters to minimize false positives. The primary ones we used are that the measured pause duration per detection is positive, as our method can lead to negative pause durations, and that the pause site is determined to within ± 100 bp (99% confidence interval); see the end of this note for the secondary filters. We determined pause site confidence using the goodness-of-fit *g* versus candidate pause positions along each replication track generated by our approach. We assumed the probability distribution of the pause site is given by ~ exp(-*g*) as is done in standard approaches and calculated the error in pause site determination as the 99% confidence interval.

We checked if our 'cut-and-align' procedure and the subsequent primary filtration worked by detecting pauses on simulated replication tracks. We represented stochasticity of a real dataset such as varying base composition between tracks, which thymidines are replaced by BrdU, location of pauses within replisome tracks, and duration of pauses. For simplicity, we did not incorporate other sources of stochasticity such as those introduced by DNAscent detect, forkSense, and variation in replisome track length.

We generated random DNA sequences 18 kb long with spatial BrdU patterns set by the probability of substitution of thymidine by BrdU $\rho$, given by

$$\rho(x) = \begin{cases} \rho_{\text{low}} + \dfrac{\rho_{\text{high}} - \rho_{\text{low}}}{1 + \exp\left(\dfrac{x - 9000}{w}\right)}, x \le x_{\text{p}} \\ \rho_{\text{low}} + \dfrac{\rho_{\text{high}} - \rho_{\text{low}}}{1 + \exp\left(\dfrac{x - 9000 + d_{\text{p}}}{w}\right)}, x > x_{\text{p}} \end{cases}$$

These tracks with coordinate $x$ represent right-moving replisomes according to our best-fit sigmoid, with the sigmoidal centre at the centre of the track, but with one pause per track whose location $x_{\text{p}}$ and duration $d_{\text{p}}$ vary between tracks. We express durations $d_{\text{p}}$ in units of length here (kb), which can be converted to units of time as in the main text using a mean synthesis rate e.g. 2 kb min$^{-1}$. The sigmoidal parameters $\rho_{\text{high}}, \rho_{\text{low}}$ and $w$ have their usual meanings of high, low, and width respectively, and we set them to 0.11, 0.65, and 3 kb respectively; these are either the best-fit values from the main text or very close to them.

We then simulated 100,000 tracks—we set each track to have a pause or not with equal probability, then for the paused tracks drew the site $x_{\text{p}}$ and duration $d_{\text{p}}$ pair uniformly from the set of points $(x, y)$ such that $0 \le x \le 18$ kb and $0 \le y \le 18$ kb $- x$. We capped pause durations because although we can detect the location of a long-lasting pause, the associated duration is harder to measure (see below). First, we detected no pauses on all the tracks without simulated pauses. This suggests a high true-negative rate and a low false-positive rate in our real dataset. On the tracks with pauses, we measured good agreement between the simulated and the detected pause location and duration (Fig. S9). Pause sites differed by -5 ± 40 bp and durations differed by 920 ± 2400 bp from the respective actual values (mean ± s.d., $n$ = 9018 tracks). However, we detected pauses only on ~18% of tracks with pauses on them. These suggest that in the real dataset our true positive rate is close to one, but our false negative rate is large as well. This was by design as we wanted to capture pauses with confidence in our real dataset which has other sources of noise not captured by this simulation. This was also because we miss pauses that occur when the amount of BrdU is low—this is discussed below.

We observed that although the average agreement between measured and observed durations was good, the difference between these on individual events became larger as durations became longer (Fig. S9B). This is because pause durations are harder to estimate on longer pauses as the BrdU runs out. If BrdU density is low across a segment of a molecule, there is no information about the progression of synthesis across this segment, as our technique relies on a mapping between BrdU density and synthesis time. This is also why pause detection is difficult when pauses occur beyond the midpoint of the sigmoid in our simulated dataset, as the characteristic step change in BrdU density associated with pause detection is small or absent when BrdU density is low (Fig. S9A).

In analysis of the actual experimental datasets, we applied five more filters to further minimize false positives: (1) the maximum of the replisome track probability as assigned by forkSense is at least 0.9, (2) the section of the track upstream of the pause site is at least 200 bp long, (3) the section of the track downstream of the pause site is at least 2000 bp long, (4) the BrdU density at the start of the replication track is at least 0.5, and (5) the track length is at least 3 kb. In the main wildtype dataset, we found that the primary filters rejected a substantial fraction of pause sites detected on replication tracks (300282 out of 332096), and the secondary filters performed further winnowing (another 14781 removed).

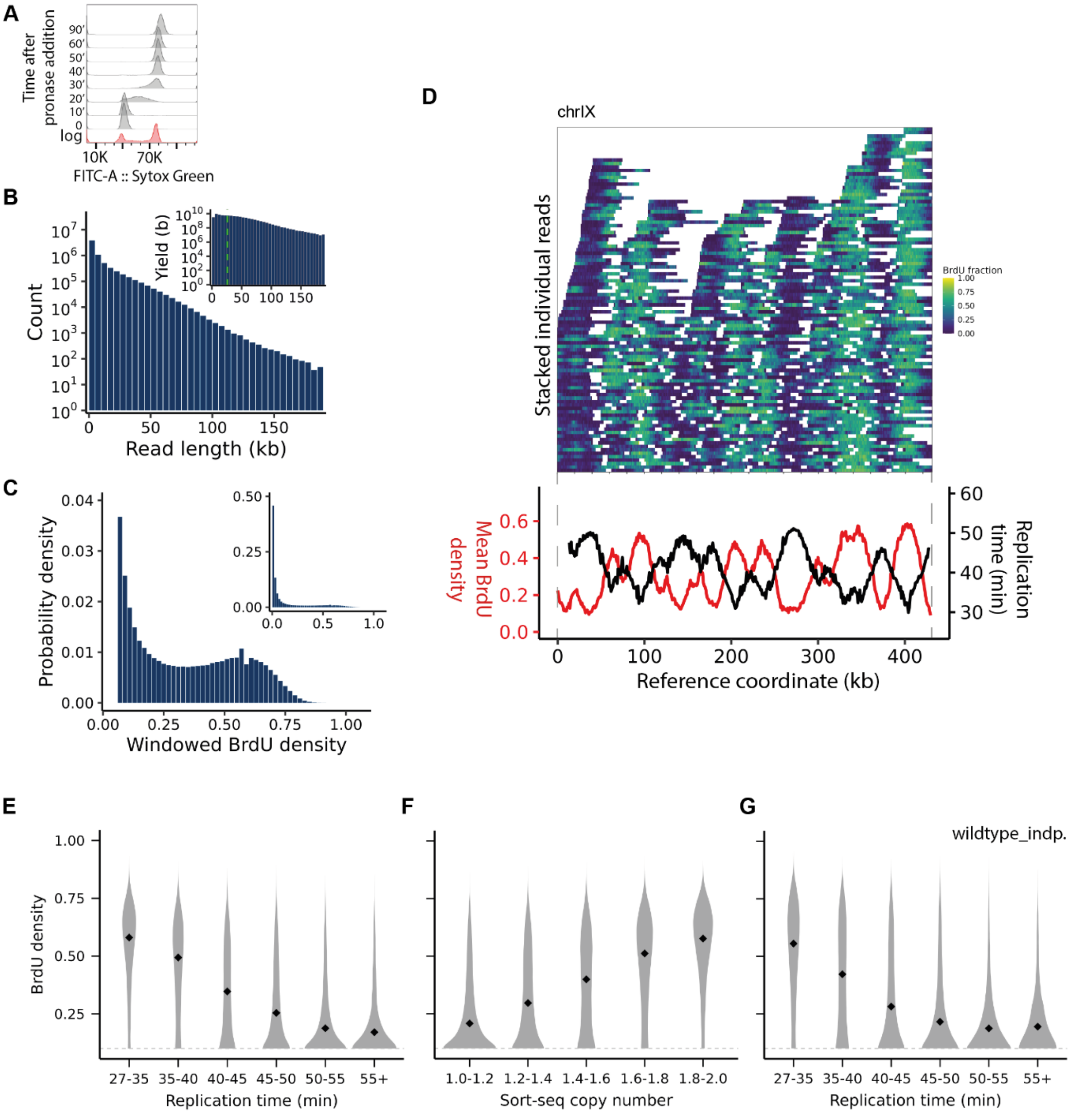


**Figure S1. Statistics of BrdU incorporation in single molecules, related to Figure 1. (A)** Flow cytometry analysis of DNA content of G1 cell cycle arrest and release into S-phase from our wildtype strain (ARY017). **(B)** Histogram of read lengths from our dataset ($n$ = 6.9 million). Inset: sequencing yield versus read length i.e. total length of reads belonging to each bin of read length. Green dashed line: N50. **(C)** Probability distribution of BrdU density measured by sectioning a randomly chosen subset of reads into non-overlapping windows of 300 thymidines along their length. Inset: same figure with density limits from 0 to 1 (calculated from $n$ = 1.8 million windows). **(D)** Top: heatmap of BrdU density versus reference coordinate across randomly chosen molecules aligning to chrIX ($n$ = 609). Heatmap of vertically and horizontally stacked molecules generated by an algorithm that sets a stacking order such that at least 5 kb separates neighbouring molecules on the same row. Bottom: BrdU density averaged across these reads and median replication time (measured in ref. (2)) versus genomic coordinates. **(E-G)** Probability distribution of BrdU density per molecule measured in 1 kb genomic windows and aggregated by corresponding ensemble measurements of median replication time **(E,G)** and copy numbers using Sort-seq **(F)**, both in the same 1 kb windows and lifted over from ref. (2) ($n$ = 38384 **(E, F)** and 74092 molecules **(G)**). Only densities above 0.1 are displayed (grey dashed line). Also see *Relation between BrdU density and replication time* from *Methods*. **(G)** is a result of analysis in this study using sequencing data from ref. (1) (labelled as 'wildtype_independent'). We used all molecules above 30 kb for the analysis in **(G)** but only a 10% randomly chosen subset in **(E)** and **(F)**.

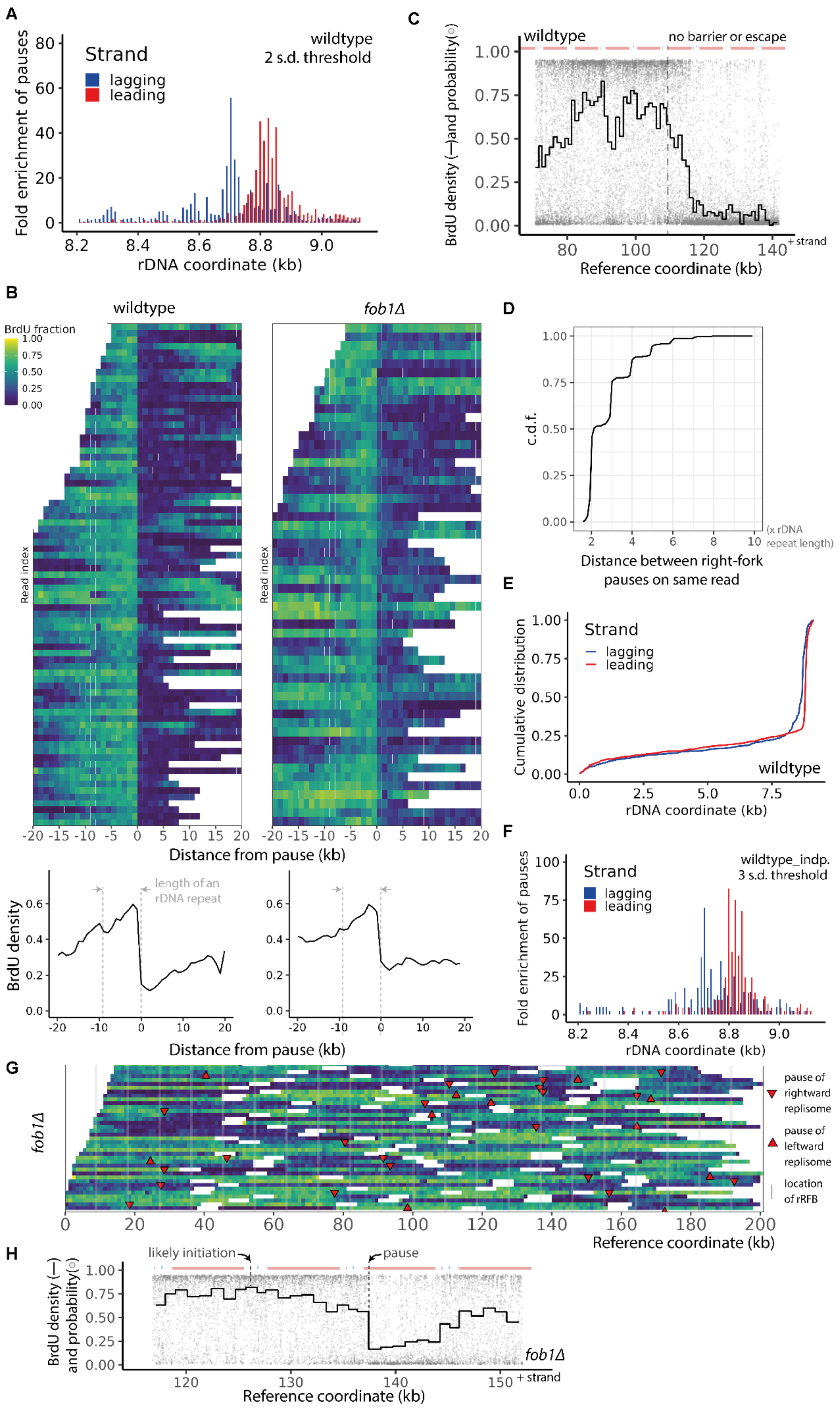

A
Fold enrichment of pauses
Strand
lagging
leading
wildtype
2 s.d. threshold
rDNA coordinate (kb)
C
BrdU density (—)and probability(○)
wildtype
no barrier or escape
Reference coordinate (kb)
+ strand
B
wildtype
fob1Δ
BrdU fraction
Read index
Distance from pause (kb)
length of an rDNA repeat
BrdU density
D
c.d.f.
(x rDNA repeat length)
Distance between right-fork pauses on same read
E
Cumulative distribution
Strand
lagging
leading
wildtype
rDNA coordinate (kb)
F
Fold enrichment of pauses
Strand
lagging
leading
wildtype_indp.
3 s.d. threshold
rDNA coordinate (kb)
G
fob1Δ
pause of rightward replisome
pause of leftward replisome
location of rRFB
Reference coordinate (kb)
H
likely initiation
pause
BrdU density (—) and probability(○)
fob1Δ
Reference coordinate (kb)
+ strand

**Figure S2. Statistics of fork pausing at the rDNA, related to Figure 1. (A)** Fold enrichment of pauses near the rRFB versus rDNA coordinate collapsed onto one rDNA repeat in wildtype cells. Analysis same as Fig. 1E except with a comparatively relaxed filtration of pauses using a 2 s.d. criterion instead of a 3 s.d. criterion (see *Methods*). Although we detect more pauses in this region (3493 vs 3041), the spatial pattern of pauses is largely unchanged. This suggests that our method is robust with respect to filtering parameters. **(B)** Top: heatmaps of a subset of molecules with detected rightward replisome pauses at the rDNA versus distance from the pause site in wildtype and *fob1Δ* cells. Molecules were aligned at their respective pause sites which were predominantly clustered at the rRFB in wildtype cells but are distributed much more diffusely throughout the rDNA repeat in *fob1Δ* cells (Figs. 1F, I). Bottom: BrdU density averaged across these molecules. Wildtype pauses are more prominent as they display a steeper drop in BrdU density. **(C)** Single-molecule view of BrdU substitution probability (grey) and average BrdU fraction over 300T windows (black) showing a rightward replisome from wildtype cells that does not pause at the barrier (black dashed line). **(D)** Cumulative distribution function (CDF) of distance between rightward replisome pauses along those molecules with multiple pause detections ($n$ = 313 distances). Distances between pauses are almost all multiples of the rDNA repeat length. **(E)** Cumulative distribution function of pause locations from Fig. 1F (data are collapsed onto one rDNA repeat) coloured by synthesis direction. Leading-strand pauses are more barrier proximal and focused. **(F)** Fold enrichment of pauses coloured by strand identity in wildtype cells from ref. (1). A 3 s.d. threshold as used in Fig. 1E was used here. **(G)** Heatmaps of BrdU density of a randomly selected subset of nascent molecules aligned to the rDNA in *fob1Δ* cells ($n$ = 131 molecules). Detected replisome pauses on rightward and leftward tracks shown with downward and upward facing arrowheads respectively. Pauses are not clustered at the rRFB unlike those in wildtype cells (Fig. 1C). **(H)** Single-molecule view of BrdU substitution probability and density showing a rightward replisome that traverses one repeat from its likely initiation site before pausing in the 35S gene of the next repeat in a *fob1Δ* cell.

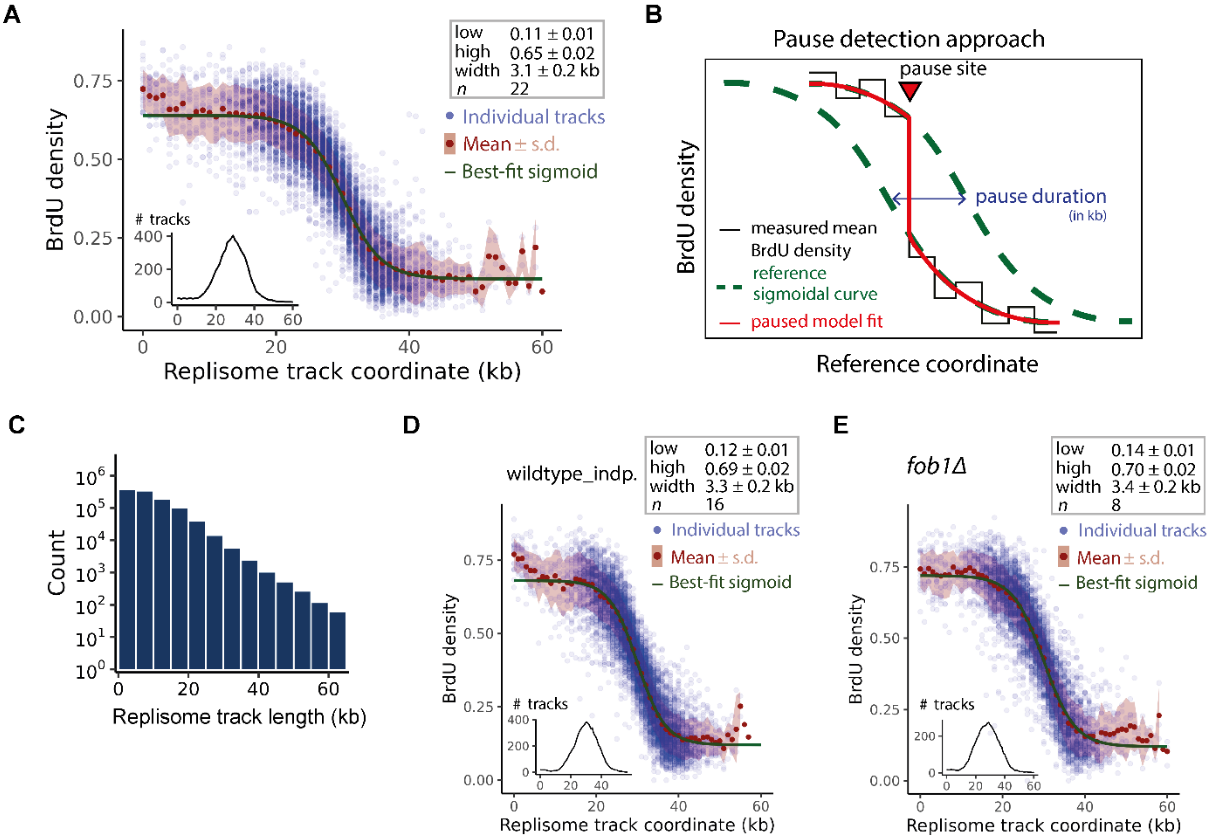


**Figure S3. Pause detection method in the non-repetitive genome, related to Figure 2. (A)** Average BrdU density per kb (blue) and sigmoidal model that best fit these tracks collectively (green) versus coordinate along a randomly-selected subset of replisome tracks ($n$ = 583, also see *Methods*). Briefly, to establish a reference BrdU density pattern versus track coordinate for pause detection, we sampled 1000 tracks, rejected those without a gradient, and shifted each horizontally to align them to a sigmoidal shape (whose center is at $x$ = 30 kb in figure). Leftward tracks were flipped to rightward before analysis. We then calculated the goodness of fit. We repeated this procedure for the same tracks but with different sets of parameters of the sigmoidal shape, and recorded a goodness of fit for each set. We then selected that parameter set that produced the best fit (green curve). We then repeated this procedure 22 times, each time drawing a new set of 1000 tracks at random, and averaged all these best-fit parameters to obtain the parameters of the best-fit sigmoid shown in the inset grey box (mean ± s.d.). Inset: Number of tracks versus coordinate. As we horizontally shifted each track in the best-fit procedure described above, not all tracks start at $x$ = 0 and end at $x$ = 60 kb. In other words, the number of tracks at each $x$ axis coordinate is different, and this is shown in the inset. **(B)** Schematic of our pause-detection method. We first convert BrdU probability at each thymidine along a replisome track to a binary state of substituted (1) or not (0) using a threshold of 0.5. We then split the track into two parts at each thymidine and fit the reference sigmoid of the grey box in **(A)** to the binary data in each part separately using just the horizontal offsets of the sigmoids as fitting parameters. We then measure the combined goodness of fit. We pick the thymidine at which the best goodness of fit occurred as the detected pause site. We then measure the pause duration as the difference between the horizontal offsets of the two sigmoids and convert it to a time assuming a fork speed of 2 kb $min^{-1}$ (3). We then apply filters to the set of detected pause sites across all replisome tracks (also see *Methods*). **(C)** Histogram of lengths of all detected replisome tracks. **(D, E)** Same analysis as **(A)** on molecules from our previous dataset (1), and molecules from *fob1Δ* cells. Parameters from **(D)** were used to find pause sites in the independent dataset using the same procedure as our primary dataset **(B)**.

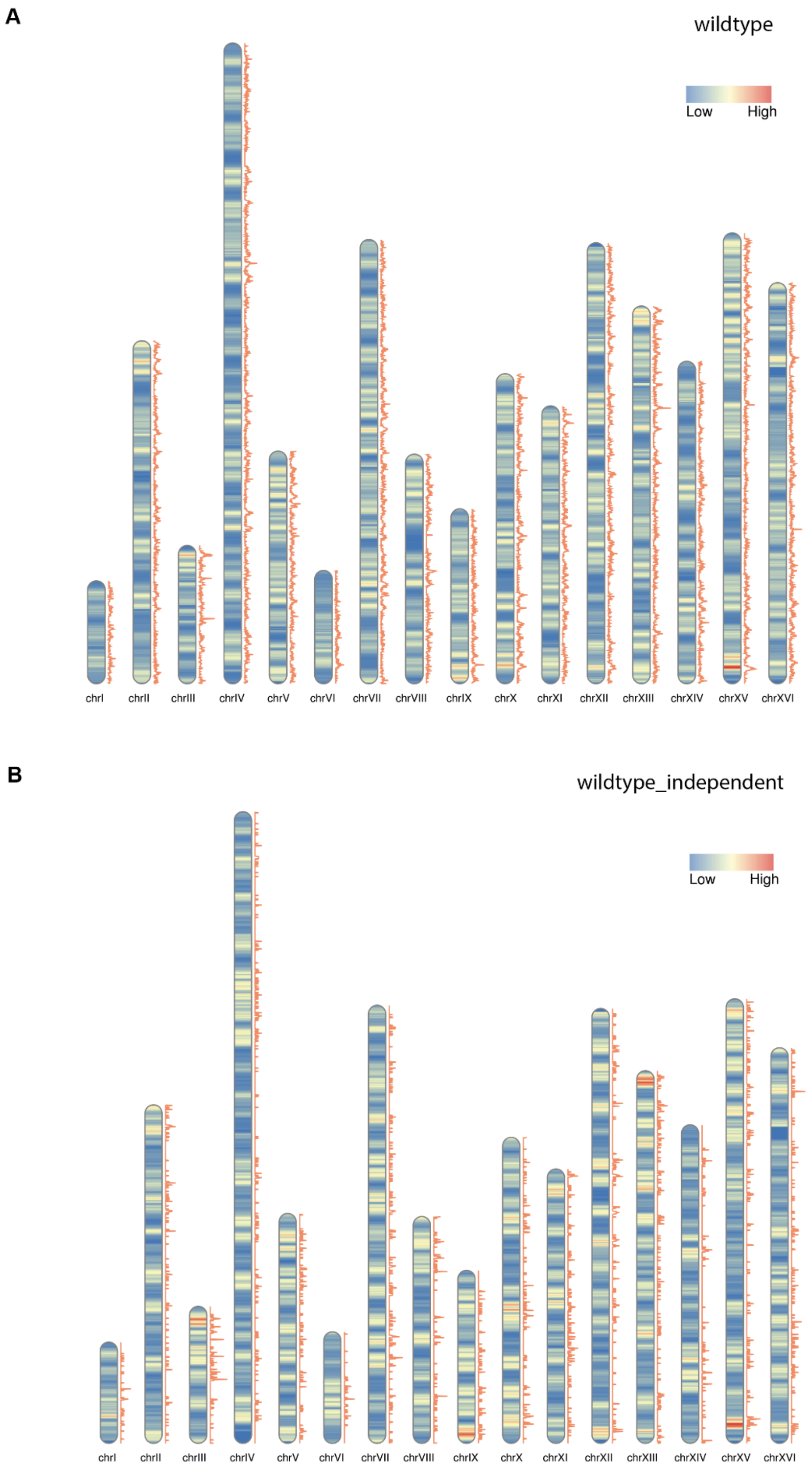


**Figure S4. Spatial profile of individual, single molecule pausing events in the non-repetitive genome, related to Figure 2.** Expected (heatmap in contigs) and observed (orange track) pause counts versus genomic coordinate in the dataset of this paper (A) and from our analysis here of molecules from our independent, previous study (1) (B).

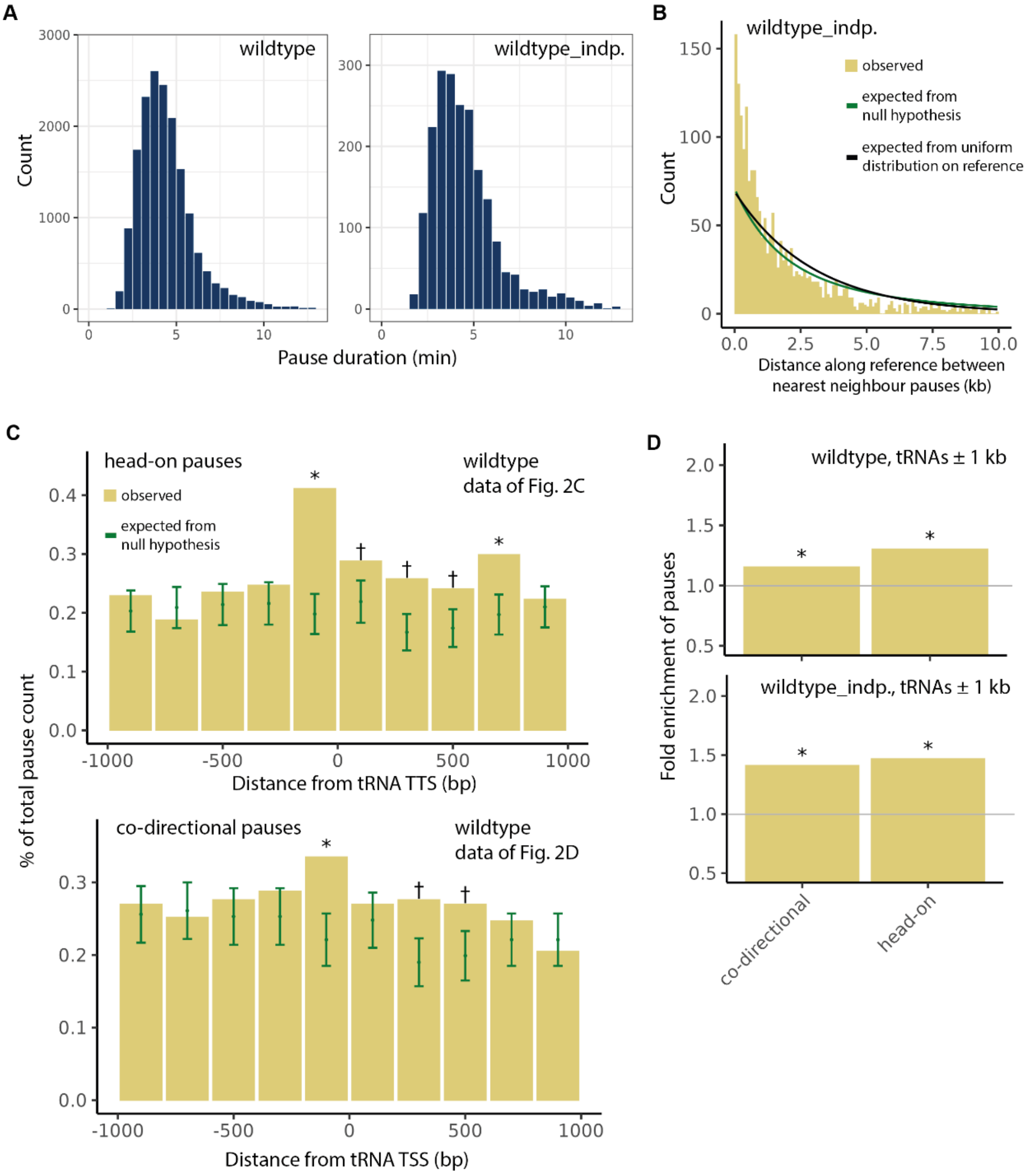


**Figure S5. Statistics of individual pause events in the non-repetitive genome and at tRNAs, related to Figure 2. (A)** Histogram of pause durations in our current dataset (left) and obtained by analysis of our previous dataset (right) (1) (*n* = 17033 and 2061 pauses respectively). Durations were calculated using the BrdU step size at the pause site and our best-fit sigmoidal model, assuming a fork speed of 2 kb $min^{-1}$ (3). **(B)** Histogram of expected and observed nearest-neighbour distances between pauses along the reference genome from our independent dataset (green and yellow respectively, *n* = 1987 distances). Black: expectation from a spatially uniform distribution of pauses. To compare with Fig. 2B, dataset size must be considered (see *Supplementary Note 1*). **(C)** Expected and observed counts versus distance of pause relative to nearest tRNA transcription termination (top) and start sites (bottom) of head-on (top) and co-directional (bottom) pauses (*: > 3 s.d, †: > 1.65 s.d. of enrichment). Figs. 2C, D show observed-to-expected ratios from here. Pause counts normalized to total pause count in the non-repetitive genome and reported as a percentage. Error bars are s.d. **(D)** Fold enrichment of pauses of either relative orientation in a 1 kb neighbourhood of tRNAs from analyses of our current study (top, *n* = 460 and 446 pauses in co-directional and head-on bins respectively) and our independent dataset (bottom, *n* = 89 and 72 pauses in co-directional and head-on bins respectively). Bin boundaries are the same as that from the top of **(C)**.

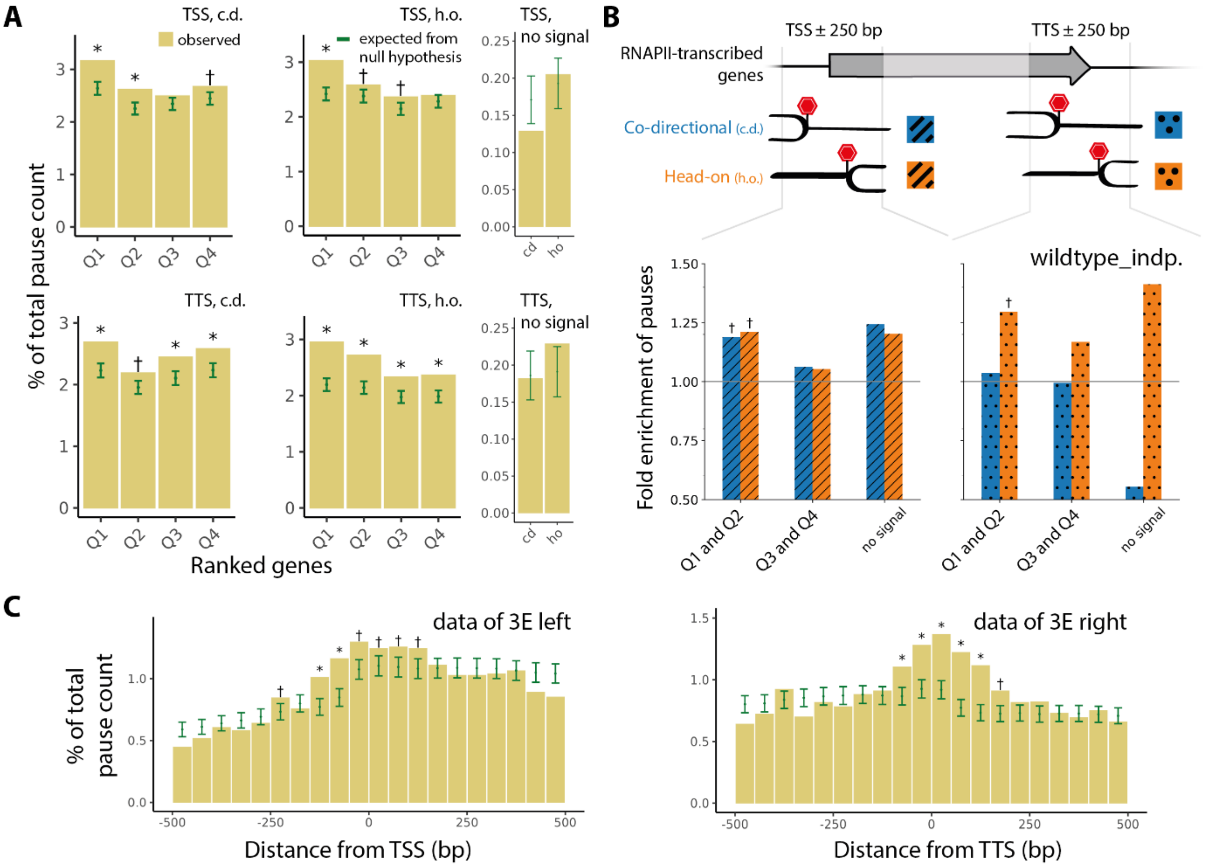


**Figure S6. Statistics of individual pause events in and near RNAPII-transcribed genes correlated with RNAPII amount, related to Figure 3. (A)** Observed and expected pause counts in 500 bp regions centred at transcription start and end sites of RNAPII-transcribed genes ordered into quartiles and one no-signal bin by RNAPII CRAC amounts. Ratio of these form data underlying top of Fig. 3A; see that figure for more details. Pause counts reported as a percentage of total pause count in the non-repetitive genome. **(B)** Same analysis as Fig. 3A but performed on molecules from our independent, previous study (1). **(C)** Observed and expected pause counts versus distance from TSS/TTS, whose ratio was taken to form Fig. 3E; see there for more details. In entire figure, * and † mean > 3 s.d or 1.65 s.d. respectively of enrichment, and errors bars are s.d.

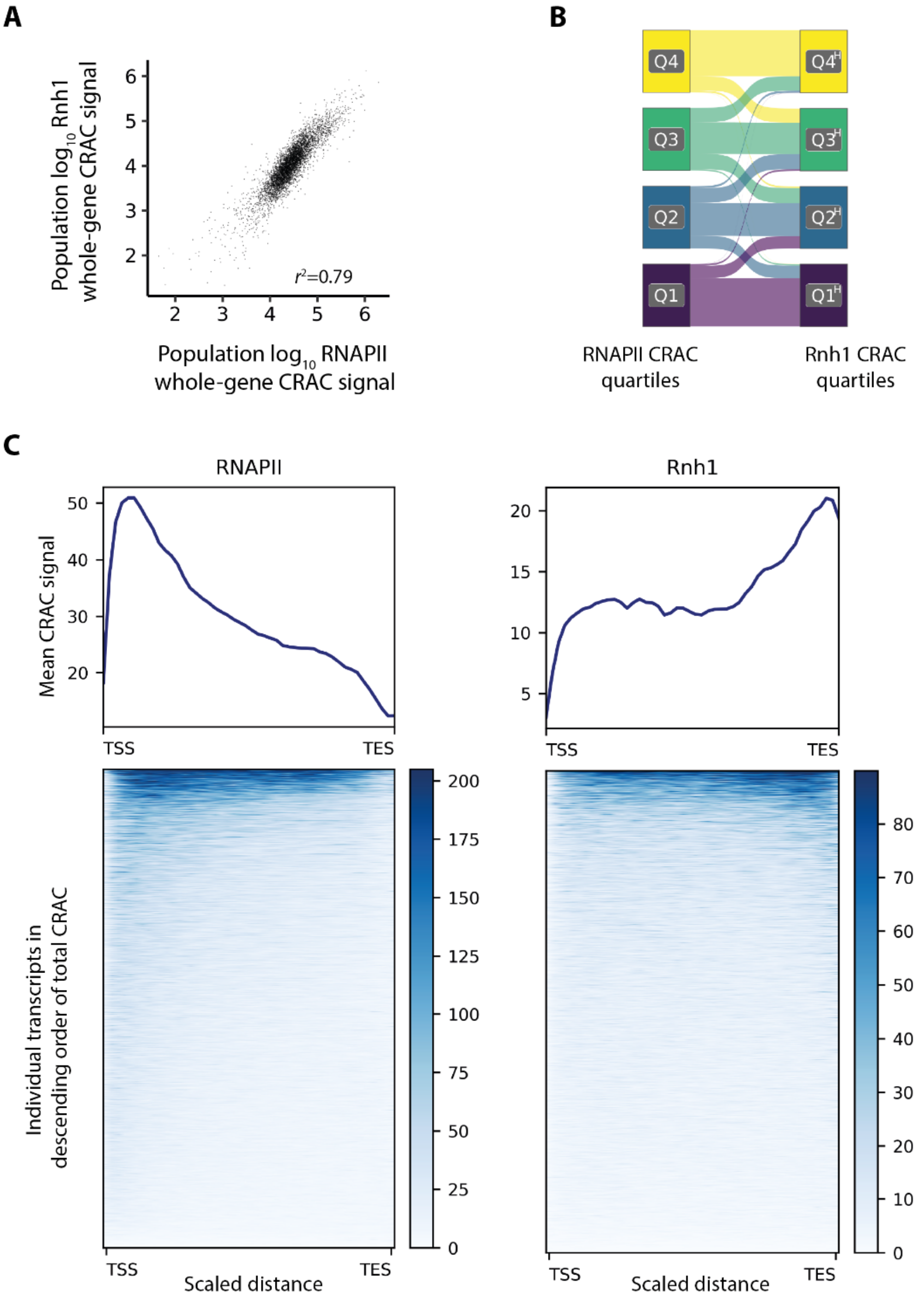


**Figure S7. Comparison between RNAPII and Rnh1 CRAC signals at RNAP-II transcribed genes, related to Figure 4.** CRAC data and transcript coordinates obtained from refs. (4), (5) respectively. **(A)** Total Rnh1 CRAC signal per gene versus total RNAPII CRAC signal per gene ($n$ = 5127 transcripts). **(B)** Sankey diagram showing relative numbers of genes that fall in the same rank in quartiles of both RNAPII and Rnh1 CRAC, and those that fall into different quartiles ($n$ = 5127 transcripts). **(C)** Top: RNAPII (left) and Rnh1 (right) CRAC signal averaged across transcripts versus scaled distance along transcript on *sacCer3* coordinates ($n$ = 5169 transcripts; also see *Visualizations* of *Methods*). Bottom: Heatmap of signal where colour is CRAC signal, each row is one transcript scaled to same length, and rows are in descending order of summed signal. Each column in heatmap was averaged and plotted on top. Our spatial profiles are similar to the median RNAPII CRAC profiles and exemplary Rnh1 CRAC profiles presented in ref. (4).

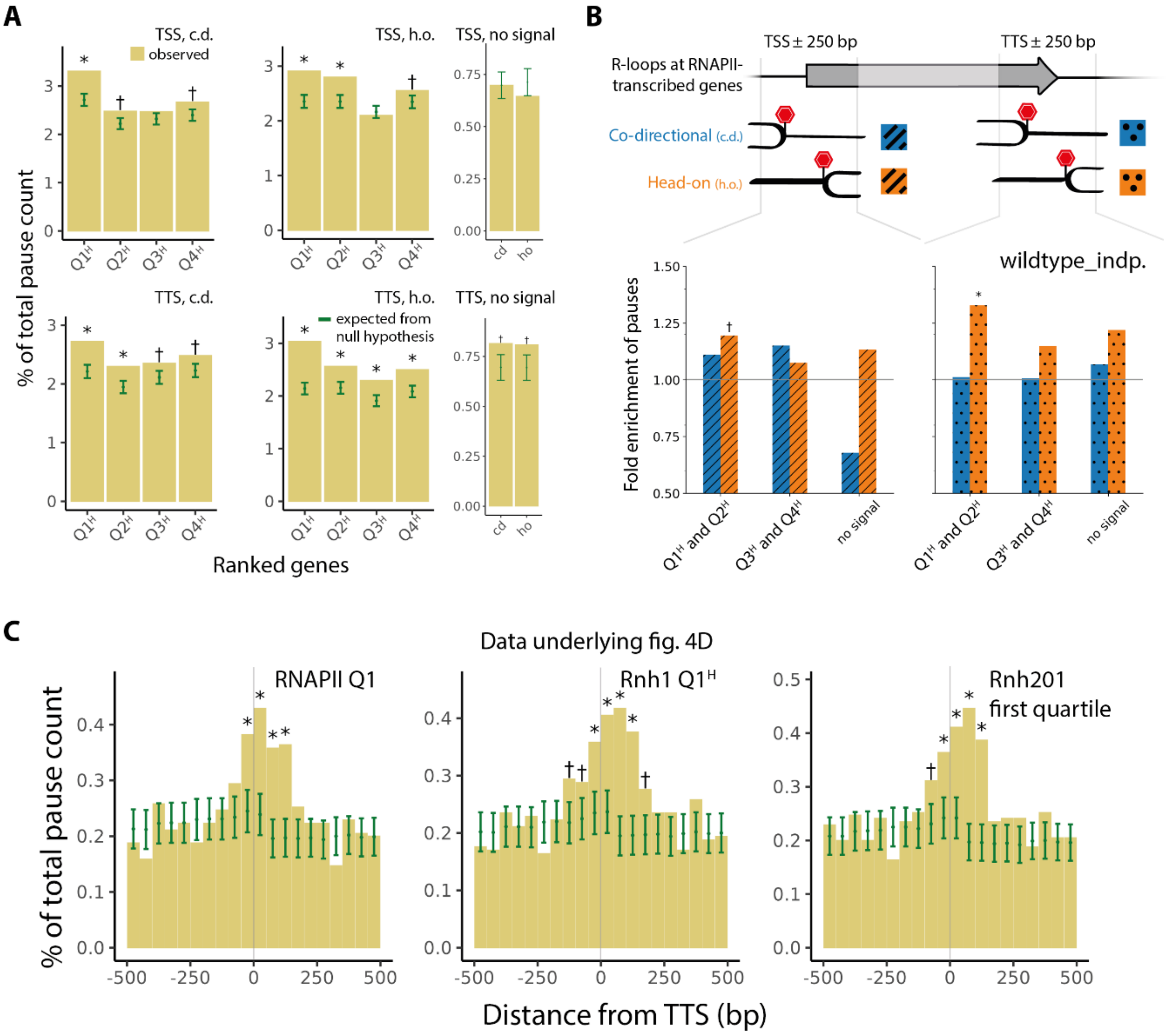


**Figure S8. Statistics of individual pause events in and near RNAPII-transcribed genes correlated with R-loop amount, related to Figure 4. (A)** Observed and expected pause counts in 500 bp regions centred at transcription start and end sites of RNAPII-transcribed genes ordered into quartiles and one no-signal bin by Rnh1 CRAC amounts. Ratio of these form data underlying top of Fig. 4A; see that figure for more details. Pause counts reported as a percentage of total pause count in the non-repetitive genome. **(B)** Same analysis as Fig. 4A but performed on molecules from our independent, previous study (1). **(C)** Observed and expected pause counts versus signed distance from TTS in the highest quartile of RNAPII, Rnh1, and Rnh201 CRAC (left to right). Ratio was taken to form Fig. 4D; see there for more details. In entire figure, * and † mean > 3 s.d or 1.65 s.d. respectively of enrichment, and errors bars are s.d.

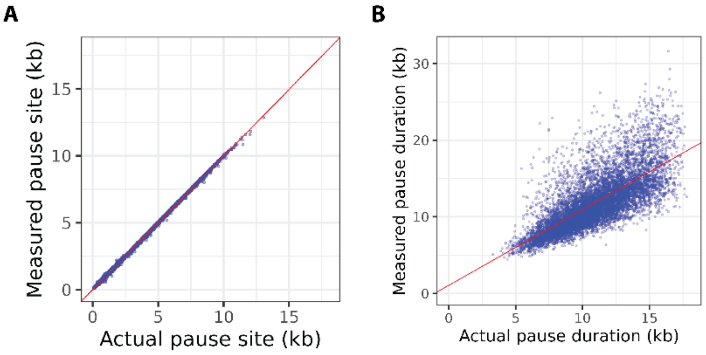


**Figure S9. Comparison between measured and actual pause sites in a simulated dataset. (A, B)** Measured versus actual pause site **(A)** and pause duration **(B)** in a dataset of simulated replisome tracks with pauses (*n* = 9018) (see Supplementary Note 2). Red: best-fit straight lines given by $y = 1.0x + 0.01$ ($R^2 = 1.0$, **(A)**) and $y = 0.99x + 1.07$ ($R^2 = 0.53$, **(B)**).

**Table S1. Strains used in this study.**

| S. no. | Strain | Genotype | Source |
|---|---|---|---|
| **1** | ARY017 | *MATa, RAD5, BUD4, leu2, ura3, trp1, ade2, his3 ChrVIII:202751::pSAC6-hENT1-tENO2-pPOP6-hsvTK-tTDH1* | This study |
| **2** | E3087 | *MATa, ade2-1, trp1-1, can1-100, leu2-3,112, his3-11,15, URA3::GPD-TK(5x), AUR1c::ADH-hENT1, RAD5+* | (6) |
| **3** | *fob1Δ* | E3087 *fob1Δ::KanMX* | This study |

**Table S2. Number of molecules analysed and pauses detected in the rDNA.**

| S. no. | Number of: | wildtype | wildtype[a] (independent, ref. (1)) | ***fob1Δ*** |
|---|---|---|---|---|
| **1** | molecules sequenced using nanopore | 6875207 | 912710 | 1789478 |
| **2** | modification-called rDNA molecules with a minimum alignment length of 30 kb (≈ 3.3 rDNA repeats) | 36039 | 5235 | 13489 |
| **3** | nascent molecules (those from row 2 with average BrdU density ≥ 0.05) | 14615 | 2344 | 6425 |
| **4** | detected replisome pauses on molecules from row 3 | 4193 | 569 | 1528 |
| | from leftward replisomes | 513 | 45 | 536 |
| | from rightward replisomes | 3680 | 524 | 992 |
| | from leading-strand synthesis | 2145 | 289 | 790 |
| | from lagging-strand synthesis | 2048 | 280 | 738 |
| **5** | detected replisome pauses from row 4 near the rRFB (within the interval 8200-9137, or equivalently -655 to 282 where 0 is the first Fob1 binding site; similar to the limits of Figs. 1E, H, S2F) | 3041 | 442 | 148 |
| | from leftward replisomes | 54 | 8 | 49 |
| | from rightward replisomes | 2987 | 434 | 99 |
| | from leading-strand synthesis | 1570 | 226 | 76 |
| | from lagging-strand synthesis | 1471 | 216 | 72 |
| **6** | molecules on which pauses from row 4 were seen | 3749 | 526 | 1338 |

[a] nanopore currents recorded in ref. (1), but every other downstream step performed in this study

**Table S3. Number of molecules analysed and pauses detected in the non-repetitive genome.**

| S. no. | Number of: | wildtype | wildtype[a] (independent, ref. (1)) |
|---|---|---|---|
| **1** | molecules sequenced using nanopore[b] | 6875207 | 912710 |
| **2** | modification-called molecules[c] | 2704457 | 469409 |
| **3** | detected replisome tracks | 1021595 | 260543 |
| **4** | replisome tracks filtered for length[d] and with successful model-fitting (see *Methods*) | 332096 | 60276 |
| **5** | pauses detected on tracks from row 4 | 17033 | 2061 |
| **6** | pauses per sequenced molecule[e] | $2.5 \times 10^{-3}$ | $2.3 \times 10^{-3}$ |
| **7** | base pairs in the *S. cerevisiae* genome[f] | 12297572 bp | |
| **8** | base pairs between pauses on genome on average[g] | 722 bp | 5967 bp |
| **9** | replisome tracks per bp of genome on average[h] | 330 | 60 |

[a] nanopore currents recorded in ref. (1), but every other downstream step performed in this study

[b] same as row 1 of Table S2

[c] only primary alignments from molecules with alignment length above 1 kb and mapping quality above 20 were analysed

[d] min. length 3 kb on a molecule with min. alignment length 30 kb not aligning to the rDNA, the mitochondrial contig, or the 2-micron plasmid

[e] row 5 divided by row 1

[f] from a W303 reference genome after excluding sequences of the rDNA, the mitochondrion, and the 2-micron plasmid, see *Methods*

[g] row 7 divided by row 5

[h] of tracks from row 4

**Table S4. Attached file showing 3 s.d. pause sites on W303 and SacCer3 coordinates.**
