## Supplementary figures and images for "Single-molecule landscape of DNA replication pausing"

### Movie 1

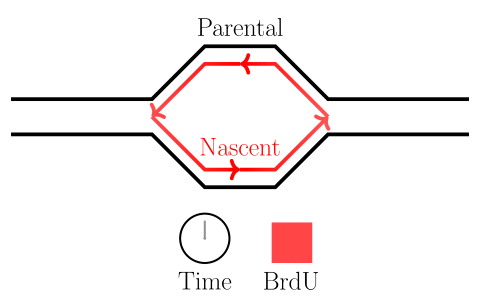

### Movie 2

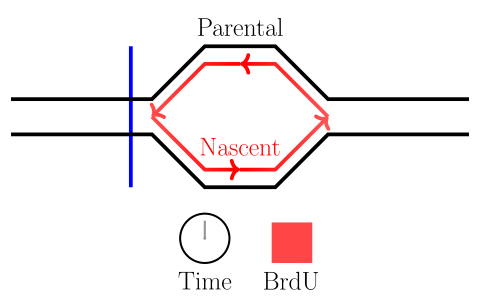
